## Supplementary material for "Dynamic genome plasticity during unisexual reproduction in the human fungal pathogen *Cryptococcus deneoformans*": Table S2.docx

**Table S2. Deletion of cell cycle regulator genes impacted diploid blastospore formation.**

| Strains | Sites dissected | Blastospores dissected | Blastospores germinated | Germination rate | Blastospores tested for ploidy | Diploid (D) | Aneuploid | Haploid (H) | Mixed H/D |
| --- | --- | --- | --- | --- | --- | --- | --- | --- | --- |
| XL280α | 10 | 49 | 47 | 96% | 20 | 18 (90%) | 2 (10%) | 0 | 0 |
| *pcl2*∆-1 | 15 | 49 | 41 | 84% | 27 | 25 (93%) | 0 | 2 (7%) | 0 |
| *pcl2*∆-2 | 15 | 40 | 22 | 55% | 17 | 17 (100%) | 0 | 0 | 0 |
| *pcl6*∆-1 | 10 | 42 | 21 | 50% | 13 | 6 (46%) | 0 | 0 | 7 (53%) |
| *pcl6*∆-2 | 10 | 42 | 28 | 67% | 17 | 5 (29%) | 0 | 2 (12%) | 10 (59%) |
| *pcl9*∆-1 | 12 | 36 | 33 | 92% | 23 | 4 (17%) | 0 | 19 (83%) | 0 |
| *pcl9*∆-2 | 12 | 31 | 30 | 97% | 24 | 6 (25%) | 0 | 18 (75%) | 0 |
| *cks1*∆-1 | 10 | 40 | 21 | 53% | 13 | 0 | 0 | 13 (100%) | 0 |
| *cks1*∆-2 | 10 | 39 | 21 | 54% | 15 | 0 | 0 | 15 (100%) | 0 |
| *cks2*∆-1 | 10 | 57 | 44 | 77% | 18 | 8 (44%) | 0 | 10 (56%) | 0 |
| *cks2*∆-2 | 10 | 53 | 40 | 75% | 18 | 10 (56%) | 0 | 8 (44%) | 0 |
| *P_GAL7_-CLB3*^*^ | 15 | 40 | 3 | 7.5% | 3 | 0 | 0 | 3 (100%) | 0 |

*: *P_GAL7_-CLB3* blastospores were dissected on YPG agar medium. No viable blastospore for *P_GAL7_-CLB3* were recovered on YPD agar medium.
