## Supplementary material for "Dynamic genome plasticity during unisexual reproduction in the human fungal pathogen *Cryptococcus deneoformans*": Table S3.docx

**Table S3. Mitotically passaged yeast cells maintained stable ploidy.**

| Strains | Blastospore ploidy | Single colony tested | Diploid (D) | Haploid (H) | Mixed H/D |
| --- | --- | --- | --- | --- | --- |
| XL280α | Diploid-1 | 8 | 8 | 0 | 0 |
|  | Diploid-2 | 8 | 8 | 0 | 0 |
|  | Diploid-3 | 8 | 8 | 0 | 0 |
|  | Diploid-4 | 8 | 8 | 0 | 0 |
| *pcl6*∆-1 | Diploid-1 | 8 | 8 | 0 | 0 |
|  | Diploid-2 | 8 | 8 | 0 | 0 |
|  | Diploid-3 | 8 | 8 | 0 | 0 |
|  | Diploid-4 | 8 | 8 | 0 | 0 |
|  | Mixed H/D-1 | 10 | 0 | 10 | 0 |
|  | Mixed H/D-2 | 10 | 3 | 6 | 1 |
|  | Mixed H/D-3 | 10 | 5 | 5 | 0 |
| *pcl6*∆-2 | Diploid-1 | 8 | 8 | 0 | 0 |
|  | Diploid-2 | 8 | 8 | 0 | 0 |
|  | Diploid-3 | 8 | 8 | 0 | 0 |
|  | Diploid-4 | 8 | 8 | 0 | 0 |
|  | Mixed H/D-1 | 10 | 1 | 9 | 0 |
|  | Mixed H/D-2 | 10 | 5 | 5 | 0 |
|  | Mixed H/D-3 | 10 | 3 | 7 | 0 |
