## Supplementary material for "Dynamic genome plasticity during unisexual reproduction in the human fungal pathogen *Cryptococcus deneoformans*": Table S4.docx

**Table S4. G1/S- and G2/M-phase population distribution profiles.** Wild type and cell cycle regulator mutants were arrested with hydroxyurea and nocodazole and population distributions were determined by FACS.

| **Samples** | **Single cells (% of total events)** | **G1/S phase (% of single cells** | **G2/M phase (% of single cells)** |  | **Percentile Color scale** |
| --- | --- | --- | --- | --- | --- |
| WT Before Arrest | 86.9 | 41 | 58.7 |  | **0** |
| WT Hydroxyurea arrest | 90.3 | 73.3 | 26.6 |  | **20** |
| WT Hydroxyurea release | 84.5 | 12.1 | 87.6 |  | **40** |
| WT Nocodazole arrest | 84.8 | 26.3 | 73 |  | **60** |
| *pcl2*∆-1 Before Arrest | 82.3 | 46.3 | 53 |  | **80** |
| *pcl2*∆-1 Hydroxyurea arrest | 92.3 | 82.8 | 16.8 |  | **100** |
| *pcl2*∆-1 Hydroxyurea release | 81.3 | 11.3 | 88.3 |  |  |
| *pcl2*∆-1 Nocodazole arrest | 89.5 | 34 | 65.5 |  |  |
| *pcl2*∆-2 Before Arrest | 80 | 39.6 | 59.9 |  |  |
| *pcl2*∆-2 Hydroxyurea arrest | 91.6 | 80.9 | 18.8 |  |  |
| *pcl2*∆-2 Hydroxyurea release | 77.2 | 8.67 | 91.1 |  |  |
| *pcl2*∆-2 Nocodazole arrest | 85.2 | 31.4 | 67.9 |  |  |
| *pcl6*∆-1 Before Arrest | 79.5 | 39 | 60.4 |  |  |
| *pcl6*∆-1 Hydroxyurea arrest | 91.2 | 77.2 | 22.4 |  |  |
| *pcl6*∆-1 Hydroxyurea release | 79.2 | 8.99 | 90.7 |  |  |
| *pcl6*∆-1 Nocodazole arrest | 85 | 24.4 | 75 |  |  |
| *pcl6*∆-2 Before Arrest | 76 | 38 | 61.3 |  |  |
| *pcl6*∆-2 Hydroxyurea arrest | 88.6 | 76 | 23.6 |  |  |
| *pcl6*∆-2 Hydroxyurea release | 81.3 | 10.1 | 89.6 |  |  |
| *pcl6*∆-2 Nocodazole arrest | 82.4 | 23.8 | 75.5 |  |  |
| *pcl9*∆-1 Before Arrest | 68.6 | 37.7 | 60.5 |  |  |
| *pcl9*∆-1 Hydroxyurea arrest | 83.9 | 72.4 | 27.4 |  |  |
| *pcl9*∆-1 Hydroxyurea release | 75.9 | 6.13 | 93.5 |  |  |
| *pcl9*∆-1 Nocodazole arrest | 77.4 | 23.2 | 76.1 |  |  |
| *pcl9*∆-2 Before Arrest | 80 | 36.9 | 62.4 |  |  |
| *pcl9*∆-2 Hydroxyurea arrest | 86.1 | 75.3 | 24.2 |  |  |
| *pcl9*∆-2 Hydroxyurea release | 78.4 | 5.83 | 93.9 |  |  |
| *pcl9*∆-2 Nocodazole arrest | 81.7 | 23.8 | 75.5 |  |  |
| *cks1*∆-1 Before Arrest | 35.1 | 6.47 | 88.3 |  |  |
| *cks1*∆-1 Hydroxyurea arrest | 24.2 | 7.77 | 81.9 |  |  |
| *cks1*∆-1 Hydroxyurea release | 12 | 14 | 82.3 |  |  |
| *cks1*∆-1 Nocodazole arrest | 42 | 20.6 | 72.9 |  |  |
| *cks1*∆-2 Before Arrest | 33.2 | 10.7 | 84.8 |  |  |
| *cks1*∆-2 Hydroxyurea arrest | 37.5 | 7.24 | 85.4 |  |  |
| *cks1*∆-2 Hydroxyurea release | 14.4 | 15.4 | 81.6 |  |  |
| *cks1*∆-2 Nocodazole arrest | 39.6 | 12.9 | 82.4 |  |  |
| *cks2*∆-1 Before Arrest | 83 | 34.2 | 65.2 |  |  |
| *cks2*∆-1 Hydroxyurea arrest | 92.6 | 76.3 | 23.6 |  |  |
| *cks2*∆-1 Hydroxyurea release | 83.3 | 11.5 | 88.3 |  |  |
| *cks2*∆-1 Nocodazole arrest | 85.8 | 28.7 | 70.7 |  |  |
| *cks2*∆-2 Before Arrest | 80.2 | 40.2 | 59.2 |  |  |
| *cks2*∆-2 Hydroxyurea arrest | 90.5 | 81.8 | 18.1 |  |  |
| *cks2*∆-2 Hydroxyurea release | 82.8 | 12.4 | 87.3 |  |  |
| *cks2*∆-2 Nocodazole arrest | 83.8 | 30.1 | 69.4 |  |  |
| WT YPG Before Arrest | 89.2 | 61.6 | 38 |  |  |
| WT YPG HU3 hours | 87.3 | 75 | 24.7 |  |  |
| WT YPG HU3 hours | 76.2 | 32.9 | 66.4 |  |  |
| WT YPG Nocodazole arrest | 87.4 | 43.3 | 55.4 |  |  |
| *P_UGE2_-CLB3-NEO* YPD Before Arrest | 34.7 | 7.11 | 89.8 |  |  |
| *P_UGE2_-CLB3-NEO* YPD Hydroxyurea arrest | 33.7 | 9.59 | 85.6 |  |  |
| *P_UGE2_-CLB3-NEO* YPD Hydroxyurea release | 18.7 | 12 | 86.4 |  |  |
| *P_UGE2_-CLB3-NEO* YPD Nocodazole arrest | 54.8 | 3.84 | 93 |  |  |
| *P_UGE2_-CLB3-NEO* YPG Before Arrest | 77.4 | 58.4 | 38.3 |  |  |
| *P_UGE2_-CLB3-NEO* YPG Hydroxyurea arrest | 81.7 | 56.8 | 41 |  |  |
| *P_UGE2_-CLB3-NEO* YPG Hydroxyurea release | 78.9 | 17.1 | 79.7 |  |  |
| *P_UGE2_-CLB3-NEO* YPG Nocodazole arrest | 83.1 | 42.2 | 53.4 |  |  |
