## Supplementary material for "Dynamic genome plasticity during unisexual reproduction in the human fungal pathogen *Cryptococcus deneoformans*": Table S5.docx

**Table S5. Strains and plasmids used in this study.**

| **Strain name** | **Genotype** | **Ploidy** | **Sources** |
| --- | --- | --- | --- |
| XL280α | WT | Haploid | [1] |
| XL280**a** | Congenic strain of XL280α | Haploid | [2] |
| MN142.6 | *MAT*α/α *ura5*∆::*NAT*/*ura5*∆::*NEO* | Diploid | [3] |
| CF1399 | *MAT*α *pcl2*∆::*NEO*-1 | Haploid | This study |
| CF1535 | *MAT*α *pcl2*∆::*NEO*-2 | Haploid | This study |
| CF1360 | *MAT*α *pcl6*∆::*NEO*-1 | Haploid | This study |
| CF1361 | *MAT*α *pcl6*∆::*NEO*-2 | Haploid | This study |
| CF1765 | *MAT*α *pcl9*∆::*NEO*-1 | Haploid | This study |
| CF1768 | *MAT*α *pcl9*∆::*NEO*-2 | Haploid | This study |
| CF1367 | *MAT*α *cks1*∆::*NEO*-1 | Haploid | This study |
| CF1414 | *MAT*α *cks1*∆::*NEO*-2 | Haploid | This study |
| CF1379 | *MAT*α *cks2*∆::*NEO*-1 | Haploid | This study |
| CF1387 | *MAT*α *cks2*∆::*NEO*-2 | Haploid | This study |
| CF1715 | *MAT*α *P_GAL7_-CLB3-NEO* | Haploid | This study |
| CF1300 | *MAT*α *SH-NURAT-NEO* | Haploid | This study |
| CF1321 | *MAT***a** *ura5*∆::*HYG* | Haploid | This study |
| CF1348 | *MAT***a** *ura5*∆::*HYG SH-NURAT-NEO* | Haploid | This study |
| CF1349 | *MAT*α *ura5*∆::*HYG SH-NURAT-NEO* | Haploid | This study |
| CF1610 | *MAT*α/α *ura5*∆::*HYG*/*ura5*∆::*HYG SH-NURAT-NEO*/*SH-NURAT-NEO*-1 | Diploid | This study |
| CF1611 | *MAT*α/α *ura5*∆::*HYG*/*ura5*∆::*HYG SH-NURAT-NEO*/*SH-NURAT-NEO*-2 | Diploid | This study |
| CF1354 | *MAT***a** *ura5*∆::*HYG SH-NURAT-NEO*/*SH-NAT-NEO*-1 | Haploid | This study |
| CF1355 | *MAT***a** *ura5*∆::*HYG SH-NURAT-NEO*/*SH-NAT-NEO*-2 | Haploid | This study |
| CF1356 | *MAT***a** *ura5*∆::*HYG SH-NURAT-NEO*/*SH-NAT-NEO*-3 | Haploid | This study |
| CF1357 | *MAT***a**/**a** *ura5*∆::*HYG*/*ura5*∆::*HYG SH-NURAT-NEO*/*SH-NAT-NEO*-4 | Diploid | This study |
| CF1358 | *MAT*α *ura5*∆::*HYG SH-NURAT-NEO*/*SH-NAT-NEO*-1 | Haploid | This study |
| CF1510 | *MAT***a** *pcl2*∆::*NEO* | Haploid | This study |
| CF1534 | *MAT***a** *pcl6*∆::*NEO* | Haploid | This study |
| CF1798 | *MAT***a** *pcl9*∆::*NEO* | Haploid | This study |
| CF1526 | *MAT***a** *cks1*∆::*NEO* | Haploid | This study |
| CF1516 | *MAT***a** *cks2*∆::*NEO* | Haploid | This study |
| CF1779 | *MAT*α *pcl2*∆::*NEO ura5*∆::*HYG SH-NURAT-NEO*-1 | Haploid | This study |
| CF1780 | *MAT*α *pcl2*∆::*NEO ura5*∆::*HYG SH-NURAT-NEO*-2 | Haploid | This study |
| CF1773 | *MAT*α *pcl6*∆::*NEO ura5*∆::*HYG SH-NURAT-NEO* | Haploid | This study |
| CF1774 | *MAT***a** *pcl6*∆::*NEO ura5*∆::*HYG SH-NURAT-NEO* | Haploid | This study |
| CF1806 | *MAT*α *pcl9*∆::*NEO ura5*∆::*HYG SH-NURAT-NEO*-1 | Haploid | This study |
| CF1807 | *MAT*α *pcl9*∆::*NEO ura5*∆::*HYG SH-NURAT-NEO*-2 | Haploid | This study |
| CF1784 | *MAT*α *cks1*∆::*NEO ura5*∆::*HYG SH-NURAT-NEO*-1 | Haploid | This study |
| CF1787 | *MAT*α *cks1*∆::*NEO ura5*∆::*HYG SH-NURAT-NEO*-2 | Haploid | This study |
| CF1770 | *MAT*α *cks2*∆::*NEO ura5*∆::*HYG SH-NURAT-NEO*-1 | Haploid | This study |
| CF1772 | *MAT*α *cks2*∆::*NEO ura5*∆::*HYG SH-NURAT-NEO*-2 | Haploid | This study |
| CF1835 | *MAT*α *P_GAL7_-CLB3-NEO ura5*∆::*HYG SH-NURAT-NEO*-1 | Haploid | This study |
| **Genotype** | **Blastospores dissected in this study*** | **Ploidy**** | |
| WT | XL280α blastospore 1 (47/49, 10 sites, site 1 5/5) | Diploid | |
|  | XL280α blastospore 2 (47/49, 10 sites, site 1 5/5) | Diploid | |
|  | XL280α blastospore 3 (47/49, 10 sites, site 1 5/5) |  | |
|  | XL280α blastospore 4 (47/49, 10 sites, site 1 5/5) |  | |
|  | XL280α blastospore 5 (47/49, 10 sites, site 1 5/5) |  | |
|  | XL280α blastospore 6 (47/49, 10 sites, site 2 6/6) | Diploid | |
|  | XL280α blastospore 7 (47/49, 10 sites, site 2 6/6) | Diploid | |
|  | XL280α blastospore 8 (47/49, 10 sites, site 2 6/6) |  | |
|  | XL280α blastospore 9 (47/49, 10 sites, site 2 6/6) |  | |
|  | XL280α blastospore 10 (47/49, 10 sites, site 2 6/6) |  | |
|  | XL280α blastospore 11 (47/49, 10 sites, site 2 6/6) |  | |
|  | XL280α blastospore 12 (47/49, 10 sites, site 3 7/7) | Diploid | |
|  | XL280α blastospore 13 (47/49, 10 sites, site 3 7/7) | Diploid | |
|  | XL280α blastospore 14 (47/49, 10 sites, site 3 7/7) |  | |
|  | XL280α blastospore 15 (47/49, 10 sites, site 3 7/7) |  | |
|  | XL280α blastospore 16 (47/49, 10 sites, site 3 7/7) |  | |
|  | XL280α blastospore 17 (47/49, 10 sites, site 3 7/7) |  | |
|  | XL280α blastospore 18 (47/49, 10 sites, site 3 7/7) |  | |
|  | XL280α blastospore 19 (47/49, 10 sites, site 4 3/3) | Diploid | |
|  | XL280α blastospore 20 (47/49, 10 sites, site 4 3/3) | Diploid | |
|  | XL280α blastospore 21 (47/49, 10 sites, site 4 3/3) |  | |
|  | XL280α blastospore 22 (47/49, 10 sites, site 5 6/6) | Aneuploid | |
|  | XL280α blastospore 23 (47/49, 10 sites, site 5 6/6) | Aneuploid | |
|  | XL280α blastospore 24 (47/49, 10 sites, site 5 6/6) |  | |
|  | XL280α blastospore 25 (47/49, 10 sites, site 5 6/6) |  | |
|  | XL280α blastospore 26 (47/49, 10 sites, site 5 6/6) |  | |
|  | XL280α blastospore 27 (47/49, 10 sites, site 5 6/6) |  | |
|  | XL280α blastospore 28 (47/49, 10 sites, site 6 5/5) | Diploid | |
|  | XL280α blastospore 29 (47/49, 10 sites, site 6 5/5) | Diploid | |
|  | XL280α blastospore 30 (47/49, 10 sites, site 6 5/5) |  | |
|  | XL280α blastospore 31 (47/49, 10 sites, site 6 5/5) |  | |
|  | XL280α blastospore 32 (47/49, 10 sites, site 6 5/5) |  | |
|  | XL280α blastospore 33 (47/49, 10 sites, site 7 4/4) | Diploid | |
|  | XL280α blastospore 34 (47/49, 10 sites, site 7 4/4) | Diploid | |
|  | XL280α blastospore 35 (47/49, 10 sites, site 7 4/4) |  | |
|  | XL280α blastospore 36 (47/49, 10 sites, site 7 4/4) |  | |
|  | XL280α blastospore 37 (47/49, 10 sites, site 8 3/4) | Diploid | |
|  | XL280α blastospore 38 (47/49, 10 sites, site 8 3/4) | Diploid | |
|  | XL280α blastospore 39 (47/49, 10 sites, site 8 3/4) |  | |
|  | XL280α blastospore 40 (47/49, 10 sites, site 9 5/5) | Diploid | |
|  | XL280α blastospore 41 (47/49, 10 sites, site 9 5/5) | Diploid | |
|  | XL280α blastospore 42 (47/49, 10 sites, site 9 5/5) |  | |
|  | XL280α blastospore 43 (47/49, 10 sites, site 9 5/5) |  | |
|  | XL280α blastospore 44 (47/49, 10 sites, site 9 5/5) |  | |
|  | XL280α blastospore 45 (47/49, 10 sites, site 10 3/4) | Diploid | |
|  | XL280α blastospore 46 (47/49, 10 sites, site 10 3/4) | Diploid | |
|  | XL280α blastospore 47 (47/49, 10 sites, site 10 3/4) |  | |
| *pcl2*∆-1 | CF1399 blastospore 1 (41/49, 15 sites, site 1 3/3) | Diploid | |
|  | CF1399 blastospore 2 (41/49, 15 sites, site 1 3/3) | Diploid | |
|  | CF1399 blastospore 3 (41/49, 15 sites, site 1 3/3) |  | |
|  | CF1399 blastospore 4 (41/49, 15 sites, site 2 4/4) | Diploid | |
|  | CF1399 blastospore 5 (41/49, 15 sites, site 2 4/4) | Diploid | |
|  | CF1399 blastospore 6 (41/49, 15 sites, site 2 4/4) |  | |
|  | CF1399 blastospore 7 (41/49, 15 sites, site 2 4/4) |  | |
|  | CF1399 blastospore 8 (41/49, 15 sites, site 3 5/5) | Diploid | |
|  | CF1399 blastospore 9 (41/49, 15 sites, site 3 5/5) | Diploid | |
|  | CF1399 blastospore 10 (41/49, 15 sites, site 3 5/5) |  | |
|  | CF1399 blastospore 11 (41/49, 15 sites, site 3 5/5) |  | |
|  | CF1399 blastospore 12 (41/49, 15 sites, site 3 5/5) |  | |
|  | CF1399 blastospore 13 (41/49, 15 sites, site 4 5/5) | Diploid | |
|  | CF1399 blastospore 14 (41/49, 15 sites, site 4 5/5) | Diploid | |
|  | CF1399 blastospore 15 (41/49, 15 sites, site 4 5/5) |  | |
|  | CF1399 blastospore 16 (41/49, 15 sites, site 4 5/5) |  | |
|  | CF1399 blastospore 17 (41/49, 15 sites, site 4 5/5) |  | |
|  | CF1399 blastospore 18 (41/49, 15 sites, site 5 2/2) | Diploid | |
|  | CF1399 blastospore 19 (41/49, 15 sites, site 5 2/2) | Diploid | |
|  | CF1399 blastospore 20 (41/49, 15 sites, site 6 3/3) | Diploid | |
|  | CF1399 blastospore 21 (41/49, 15 sites, site 6 3/3) | Diploid | |
|  | CF1399 blastospore 22 (41/49, 15 sites, site 6 3/3) |  | |
|  | CF1399 blastospore 23 (41/49, 15 sites, site 7 1/3) | Diploid | |
|  | CF1399 blastospore 24 (41/49, 15 sites, site 9 3/3) | Diploid | |
|  | CF1399 blastospore 25 (41/49, 15 sites, site 9 3/3) | Diploid | |
|  | CF1399 blastospore 26 (41/49, 15 sites, site 9 3/3) |  | |
|  | CF1399 blastospore 27 (41/49, 15 sites, site 10 2/2) | Diploid | |
|  | CF1399 blastospore 28 (41/49, 15 sites, site 10 2/2) | Diploid | |
|  | CF1399 blastospore 29 (41/49, 15 sites, site 11 4/4) | Diploid | |
|  | CF1399 blastospore 30 (41/49, 15 sites, site 11 4/4) | Diploid | |
|  | CF1399 blastospore 31 (41/49, 15 sites, site 11 4/4) |  | |
|  | CF1399 blastospore 32 (41/49, 15 sites, site 11 4/4) |  | |
|  | CF1399 blastospore 33 (41/49, 15 sites, site 12 2/2) | Diploid | |
|  | CF1399 blastospore 34 (41/49, 15 sites, site 12 2/2) | Diploid | |
|  | CF1399 blastospore 35 (41/49, 15 sites, site 13 2/3) | Diploid | |
|  | CF1399 blastospore 36 (41/49, 15 sites, site 13 2/3) | Diploid | |
|  | CF1399 blastospore 37 (41/49, 15 sites, site 14 2/4) | Diploid | |
|  | CF1399 blastospore 38 (41/49, 15 sites, site 14 2/4) | Diploid | |
|  | CF1399 blastospore 39 (41/49, 15 sites, site 15 3/3) | Diploid | |
|  | CF1399 blastospore 40 (41/49, 15 sites, site 15 3/3) | Diploid | |
|  | CF1399 blastospore 41 (41/49, 15 sites, site 15 3/3) |  | |
| *pcl2*∆-2 | CF1535 blastospore 1 (22/40, 15 spots, spot2 4/5) | Diploid | |
|  | CF1535 blastospore 2 (22/40, 15 spots, spot2 4/5) |  | |
|  | CF1535 blastospore 3 (22/40, 15 spots, spot2 4/5) |  | |
|  | CF1535 blastospore 4 (22/40, 15 spots, spot2 4/5) | Diploid | |
|  | CF1535 blastospore 5 (22/40, 15 spots, spot3 1/3) | Diploid | |
|  | CF1535 blastospore 6 (22/40, 15 spots, spot5 1/3) | Diploid | |
|  | CF1535 blastospore 7 (22/40, 15 spots, spot6 2/3) | Diploid | |
|  | CF1535 blastospore 8 (22/40, 15 spots, spot6 2/3) | Diploid | |
|  | CF1535 blastospore 9 (22/40, 15 spots, spot7 4/4) | Diploid | |
|  | CF1535 blastospore 10 (22/40, 15 spots, spot7 4/4) |  | |
|  | CF1535 blastospore 11 (22/40, 15 spots, spot7 4/4) |  | |
|  | CF1535 blastospore 12 (22/40, 15 spots, spot7 4/4) | Diploid | |
|  | CF1535 blastospore 13 (22/40, 15 spots, spot9 1/3) | Diploid | |
|  | CF1535 blastospore 14 (22/40, 15 spots, spot11 3/3) | Diploid | |
|  | CF1535 blastospore 15 (22/40, 15 spots, spot11 3/3) |  | |
|  | CF1535 blastospore 16 (22/40, 15 spots, spot11 3/3) | Diploid | |
|  | CF1535 blastospore 17 (22/40, 15 spots, spot13 2/2) | Diploid | |
|  | CF1535 blastospore 18 (22/40, 15 spots, spot13 2/2) | Diploid | |
|  | CF1535 blastospore 19 (22/40, 15 spots, spot14 2/3) | Diploid | |
|  | CF1535 blastospore 20 (22/40, 15 spots, spot14 2/3) | Diploid | |
|  | CF1535 blastospore 21 (22/40, 15 spots, spot15 2/2) | Diploid | |
|  | CF1535 blastospore 22 (22/40, 15 spots, spot15 2/2) | Diploid | |
| *pcl6*∆-1 | CF1360 blastospore 1 (21/42, 10 sites, site 1 2/4) | Haploid/Diploid mix | |
|  | CF1360 blastospore 2 (21/42, 10 sites, site 1 2/4) | Haploid/Diploid mix | |
|  | CF1360 blastospore 3 (21/42, 10 sites, site 2 4/4) | Diploid | |
|  | CF1360 blastospore 4 (21/42, 10 sites, site 2 4/4) | Diploid | |
|  | CF1360 blastospore 5 (21/42, 10 sites, site 2 4/4) |  | |
|  | CF1360 blastospore 6 (21/42, 10 sites, site 2 4/4) |  | |
|  | CF1360 blastospore 7 (21/42, 10 sites, site 3 2/4) | Diploid | |
|  | CF1360 blastospore 8 (21/42, 10 sites, site 3 2/4) | Diploid | |
|  | CF1360 blastospore 9 (21/42, 10 sites, site 5 5/5) | Haploid/Diploid mix | |
|  | CF1360 blastospore 10 (21/42, 10 sites, site 5 5/5) | Haploid/Diploid mix | |
|  | CF1360 blastospore 11 (21/42, 10 sites, site 5 5/5) |  | |
|  | CF1360 blastospore 12 (21/42, 10 sites, site 5 5/5) |  | |
|  | CF1360 blastospore 13 (21/42, 10 sites, site 5 5/5) |  | |
|  | CF1360 blastospore 14 (21/42, 10 sites, site 7 5/5) | Haploid/Diploid mix | |
|  | CF1360 blastospore 15 (21/42, 10 sites, site 7 5/5) | Haploid/Diploid mix | |
|  | CF1360 blastospore 16 (21/42, 10 sites, site 7 5/5) |  | |
|  | CF1360 blastospore 17 (21/42, 10 sites, site 7 5/5) |  | |
|  | CF1360 blastospore 18 (21/42, 10 sites, site 7 5/5) |  | |
|  | CF1360 blastospore 19 (21/42, 10 sites, site 8 1/4) | Haploid/Diploid mix | |
|  | CF1360 blastospore 20 (21/42, 10 sites, site 10 2/3) | Diploid | |
|  | CF1360 blastospore 21 (21/42, 10 sites, site 10 2/3) | Diploid | |
| *pcl6*∆-2 | CF1361 blastospore 1 (28/42, 10 sites, site 1 2/5) | Haploid/Diploid mix | |
|  | CF1361 blastospore 2 (28/42, 10 sites, site 1 2/5) | Haploid/Diploid mix | |
|  | CF1361 blastospore 3 (28/42, 10 sites, site 3 2/4) | Haploid | |
|  | CF1361 blastospore 4 (28/42, 10 sites, site 3 2/4) | Haploid | |
|  | CF1361 blastospore 5 (28/42, 10 sites, site 4 5/5) | Haploid/Diploid mix | |
|  | CF1361 blastospore 6 (28/42, 10 sites, site 4 5/5) | Haploid/Diploid mix | |
|  | CF1361 blastospore 7 (28/42, 10 sites, site 4 5/5) |  | |
|  | CF1361 blastospore 8 (28/42, 10 sites, site 4 5/5) |  | |
|  | CF1361 blastospore 9 (28/42, 10 sites, site 4 5/5) |  | |
|  | CF1361 blastospore 10 (28/42, 10 sites, site 5 3/4) | Diploid | |
|  | CF1361 blastospore 11 (28/42, 10 sites, site 5 3/4) | Diploid | |
|  | CF1361 blastospore 12 (28/42, 10 sites, site 5 3/4) |  | |
|  | CF1361 blastospore 13 (28/42, 10 sites, site 6 4/4) | Haploid/Diploid mix | |
|  | CF1361 blastospore 14 (28/42, 10 sites, site 6 4/4) | Haploid/Diploid mix | |
|  | CF1361 blastospore 15 (28/42, 10 sites, site 6 4/4) |  | |
|  | CF1361 blastospore 16 (28/42, 10 sites, site 6 4/4) |  | |
|  | CF1361 blastospore 17 (28/42, 10 sites, site 7 4/4) | Haploid/Diploid mix | |
|  | CF1361 blastospore 18 (28/42, 10 sites, site 7 4/4) | Haploid/Diploid mix | |
|  | CF1361 blastospore 19 (28/42, 10 sites, site 7 4/4) |  | |
|  | CF1361 blastospore 20 (28/42, 10 sites, site 7 4/4) |  | |
|  | CF1361 blastospore 21 (28/42, 10 sites, site 8 1/7) | Diploid | |
|  | CF1361 blastospore 22 (28/42, 10 sites, site 9 4/5) | Haploid/Diploid mix | |
|  | CF1361 blastospore 23 (28/42, 10 sites, site 9 4/5) | Haploid/Diploid mix | |
|  | CF1361 blastospore 24 (28/42, 10 sites, site 9 4/5) |  | |
|  | CF1361 blastospore 25 (28/42, 10 sites, site 9 4/5) |  | |
|  | CF1361 blastospore 26 (28/42, 10 sites, site 10 3/3) | Diploid | |
|  | CF1361 blastospore 27 (28/42, 10 sites, site 10 3/3) | Diploid | |
|  | CF1361 blastospore 28 (28/42, 10 sites, site 10 3/3) |  | |
| *pcl9*∆-1 | CF1765 blastospore 1 (33/36, 12 spots, spot1 4/4) | Diploid | |
|  | CF1765 blastospore 2 (33/36, 12 spots, spot1 4/4) |  | |
|  | CF1765 blastospore 3 (33/36, 12 spots, spot1 4/4) |  | |
|  | CF1765 blastospore 4 (33/36, 12 spots, spot1 4/4) | Diploid | |
|  | CF1765 blastospore 5 (33/36, 12 spots, spot2 2/3) | Haploid | |
|  | CF1765 blastospore 6 (33/36, 12 spots, spot2 2/3) | Haploid | |
|  | CF1765 blastospore 7 (33/36, 12 spots, spot3 2/2) | Haploid | |
|  | CF1765 blastospore 8 (33/36, 12 spots, spot3 2/2) | Haploid | |
|  | CF1765 blastospore 9 (33/36, 12 spots, spot4 2/2) | Haploid | |
|  | CF1765 blastospore 10 (33/36, 12 spots, spot4 2/2) | Haploid | |
|  | CF1765 blastospore 11 (33/36, 12 spots, spot5 3/3) | Haploid | |
|  | CF1765 blastospore 12 (33/36, 12 spots, spot5 3/3) |  | |
|  | CF1765 blastospore 13 (33/36, 12 spots, spot5 3/3) | Haploid | |
|  | CF1765 blastospore 14 (33/36, 12 spots, spot6 3/3) | Haploid | |
|  | CF1765 blastospore 15 (33/36, 12 spots, spot6 3/3) |  | |
|  | CF1765 blastospore 16 (33/36, 12 spots, spot6 3/3) | Haploid | |
|  | CF1765 blastospore 17 (33/36, 12 spots, spot7 2/3) | Haploid | |
|  | CF1765 blastospore 18 (33/36, 12 spots, spot7 2/3) | Haploid | |
|  | CF1765 blastospore 19 (33/36, 12 spots, spot8 4/4) | Haploid | |
|  | CF1765 blastospore 20 (33/36, 12 spots, spot8 4/4) |  | |
|  | CF1765 blastospore 21 (33/36, 12 spots, spot8 4/4) |  | |
|  | CF1765 blastospore 22 (33/36, 12 spots, spot8 4/4) | Haploid | |
|  | CF1765 blastospore 23 (33/36, 12 spots, spot9 1/2) | Haploid | |
|  | CF1765 blastospore 24 (33/36, 12 spots, spot10 4/4) | Diploid | |
|  | CF1765 blastospore 25 (33/36, 12 spots, spot10 4/4) |  | |
|  | CF1765 blastospore 26 (33/36, 12 spots, spot10 4/4) |  | |
|  | CF1765 blastospore 27 (33/36, 12 spots, spot10 4/4) | Diploid | |
|  | CF1765 blastospore 28 (33/36, 12 spots, spot 11 3/3) | Haploid | |
|  | CF1765 blastospore 29 (33/36, 12 spots, spot 11 3/3) |  | |
|  | CF1765 blastospore 30 (33/36, 12 spots, spot 11 3/3) | Haploid | |
|  | CF1765 blastospore 31 (33/36, 12 spots, spot 12 3/3) | Haploid | |
|  | CF1765 blastospore 32 (33/36, 12 spots, spot 12 3/3) |  | |
|  | CF1765 blastospore 33 (33/36, 12 spots, spot 12 3/3) | Haploid | |
| *pcl9*∆-2 | CF1768 blastospore 1 (30/31, 12 spots, spot1 2/2) | Haploid | |
|  | CF1768 blastospore 2 (30/31, 12 spots, spot1 2/2) | Haploid | |
|  | CF1768 blastospore 3 (30/31, 12 spots, spot2 2/2) | Haploid | |
|  | CF1768 blastospore 4 (30/31, 12 spots, spot2 2/2) | Haploid | |
|  | CF1768 blastospore 5 (30/31, 12 spots, spot3 3/3) | Haploid | |
|  | CF1768 blastospore 6 (30/31, 12 spots, spot3 3/3) |  | |
|  | CF1768 blastospore 7 (30/31, 12 spots, spot3 3/3) | Haploid | |
|  | CF1768 blastospore 8 (30/31, 12 spots, spot4 3/3) | Diploid | |
|  | CF1768 blastospore 9 (30/31, 12 spots, spot4 3/3) |  | |
|  | CF1768 blastospore 10 (30/31, 12 spots, spot4 3/3) | Diploid | |
|  | CF1768 blastospore 11 (30/31, 12 spots, spot5 2/3) | Haploid | |
|  | CF1768 blastospore 12 (30/31, 12 spots, spot5 2/3) | Haploid | |
|  | CF1768 blastospore 13 (30/31, 12 spots, spot6 3/3) | Haploid | |
|  | CF1768 blastospore 14 (30/31, 12 spots, spot6 3/3) |  | |
|  | CF1768 blastospore 15 (30/31, 12 spots, spot6 3/3) | Haploid | |
|  | CF1768 blastospore 16 (30/31, 12 spots, spot7 2/2) | Diploid | |
|  | CF1768 blastospore 17 (30/31, 12 spots, spot7 2/2) | Diploid | |
|  | CF1768 blastospore 18 (30/31, 12 spots, spot8 3/3) | Haploid | |
|  | CF1768 blastospore 19 (30/31, 12 spots, spot8 3/3) |  | |
|  | CF1768 blastospore 20 (30/31, 12 spots, spot8 3/3) | Haploid | |
|  | CF1768 blastospore 21 (30/31, 12 spots, spot9 3/3) | Diploid | |
|  | CF1768 blastospore 22 (30/31, 12 spots, spot9 3/3) |  | |
|  | CF1768 blastospore 23 (30/31, 12 spots, spot9 3/3) | Diploid | |
|  | CF1768 blastospore 24 (30/31, 12 spots, spot10 3/3) | Haploid | |
|  | CF1768 blastospore 25 (30/31, 12 spots, spot10 3/3) |  | |
|  | CF1768 blastospore 26 (30/31, 12 spots, spot10 3/3) | Haploid | |
|  | CF1768 blastospore 27 (30/31, 12 spots, spot11 2/2) | Haploid | |
|  | CF1768 blastospore 28 (30/31, 12 spots, spot11 2/2) | Haploid | |
|  | CF1768 blastospore 29 (30/31, 12 spots, spot12 2/2) | Haploid | |
|  | CF1768 blastospore 30 (30/31, 12 spots, spot12 2/2) | Haploid | |
| *cks1*∆-1 | CF1367 blastospore 1 (21/40, 10 sites, site 3 3/4) | Haploid | |
|  | CF1367 blastospore 2 (21/40, 10 sites, site 3 3/4) | Haploid | |
|  | CF1367 blastospore 3 (21/40, 10 sites, site 3 3/4) |  | |
|  | CF1367 blastospore 4 (21/40, 10 sites, site 5 1/4) | Haploid | |
|  | CF1367 blastospore 5 (21/40, 10 sites, site 6 4/4) | Haploid | |
|  | CF1367 blastospore 6 (21/40, 10 sites, site 6 4/4) | Haploid | |
|  | CF1367 blastospore 7 (21/40, 10 sites, site 6 4/4) |  | |
|  | CF1367 blastospore 8 (21/40, 10 sites, site 6 4/4) |  | |
|  | CF1367 blastospore 9 (21/40, 10 sites, site 7 3/3) | Haploid | |
|  | CF1367 blastospore 10 (21/40, 10 sites, site 7 3/3) | Haploid | |
|  | CF1367 blastospore 11 (21/40, 10 sites, site 7 3/3) |  | |
|  | CF1367 blastospore 12 (21/40, 10 sites, site 8 4/4) | Haploid | |
|  | CF1367 blastospore 13 (21/40, 10 sites, site 8 4/4) | Haploid | |
|  | CF1367 blastospore 14 (21/40, 10 sites, site 8 4/4) |  | |
|  | CF1367 blastospore 15 (21/40, 10 sites, site 8 4/4) |  | |
|  | CF1367 blastospore 16 (21/40, 10 sites, site 9 2/4) | Haploid | |
|  | CF1367 blastospore 17 (21/40, 10 sites, site 9 2/4) | Haploid | |
|  | CF1367 blastospore 18 (21/40, 10 sites, site 10 4/5) | Haploid | |
|  | CF1367 blastospore 19 (21/40, 10 sites, site 10 4/5) | Haploid | |
|  | CF1367 blastospore 20 (21/40, 10 sites, site 10 4/5) |  | |
|  | CF1367 blastospore 21 (21/40, 10 sites, site 10 4/5) |  | |
| *cks1*∆-2 | CF1414 blastospore 1 (21/39, 11 sites, site1 5/5) | Haploid | |
|  | CF1414 blastospore 2 (21/39, 11 sites, site1 5/5) | Haploid | |
|  | CF1414 blastospore 3 (21/39, 11 sites, site1 5/5) |  | |
|  | CF1414 blastospore 4 (21/39, 11 sites, site1 5/5) |  | |
|  | CF1414 blastospore 5 (21/39, 11 sites, site1 5/5) |  | |
|  | CF1414 blastospore 6 (21/39, 11 sites, site2 3/4) | Haploid | |
|  | CF1414 blastospore 7 (21/39, 11 sites, site2 3/4) | Haploid | |
|  | CF1414 blastospore 8 (21/39, 11 sites, site2 3/4) |  | |
|  | CF1414 blastospore 9 (21/39, 11 sites, site3 3/4) | Haploid | |
|  | CF1414 blastospore 10 (21/39, 11 sites, site3 3/4) | Haploid | |
|  | CF1414 blastospore 11 (21/39, 11 sites, site3 3/4) |  | |
|  | CF1414 blastospore 12 (21/39, 11 sites, site6 1/3) | Haploid | |
|  | CF1414 blastospore 13 (21/39, 11 sites, site7 1/2) | Haploid | |
|  | CF1414 blastospore 14 (21/39, 11 sites, site8 3/4) | Haploid | |
|  | CF1414 blastospore 15 (21/39, 11 sites, site8 3/4) | Haploid | |
|  | CF1414 blastospore 16 (21/39, 11 sites, site8 3/4) |  | |
|  | CF1414 blastospore 17 (21/39, 11 sites, site9 2/3) | Haploid | |
|  | CF1414 blastospore 18 (21/39, 11 sites, site9 2/3) | Haploid | |
|  | CF1414 blastospore 19 (21/39, 11 sites, site10 2/4) | Haploid | |
|  | CF1414 blastospore 20 (21/39, 11 sites, site10 2/4) | Haploid | |
|  | CF1414 blastospore 21 (21/39, 11 sites, site11 1/2) | Haploid | |
| *cks2*∆-1 | CF1379 blastospore 1 (44/57, 10 sites, site 1 7/8) | Diploid | |
|  | CF1379 blastospore 2 (44/57, 10 sites, site 1 7/8) | Diploid | |
|  | CF1379 blastospore 3 (44/57, 10 sites, site 1 7/8) |  | |
|  | CF1379 blastospore 4 (44/57, 10 sites, site 1 7/8) |  | |
|  | CF1379 blastospore 5 (44/57, 10 sites, site 1 7/8) |  | |
|  | CF1379 blastospore 6 (44/57, 10 sites, site 1 7/8) |  | |
|  | CF1379 blastospore 7 (44/57, 10 sites, site 1 7/8) |  | |
|  | CF1379 blastospore 8 (44/57, 10 sites, site 2 2/6) | Diploid | |
|  | CF1379 blastospore 9 (44/57, 10 sites, site 2 2/6) | Diploid | |
|  | CF1379 blastospore 10 (44/57, 10 sites, site 4 3/5) | Diploid | |
|  | CF1379 blastospore 11 (44/57, 10 sites, site 4 3/5) | Diploid | |
|  | CF1379 blastospore 12 (44/57, 10 sites, site 4 3/5) |  | |
|  | CF1379 blastospore 13 (44/57, 10 sites, site 5 7/7) | Haploid | |
|  | CF1379 blastospore 14 (44/57, 10 sites, site 5 7/7) | Haploid | |
|  | CF1379 blastospore 15 (44/57, 10 sites, site 5 7/7) |  | |
|  | CF1379 blastospore 16 (44/57, 10 sites, site 5 7/7) |  | |
|  | CF1379 blastospore 17 (44/57, 10 sites, site 5 7/7) |  | |
|  | CF1379 blastospore 18 (44/57, 10 sites, site 5 7/7) |  | |
|  | CF1379 blastospore 19 (44/57, 10 sites, site 5 7/7) |  | |
|  | CF1379 blastospore 20 (44/57, 10 sites, site 6 4/5) | Haploid | |
|  | CF1379 blastospore 21 (44/57, 10 sites, site 6 4/5) | Haploid | |
|  | CF1379 blastospore 22 (44/57, 10 sites, site 6 4/5) |  | |
|  | CF1379 blastospore 23 (44/57, 10 sites, site 6 4/5) |  | |
|  | CF1379 blastospore 24 (44/57, 10 sites, site 7 4/5) | Haploid | |
|  | CF1379 blastospore 25 (44/57, 10 sites, site 7 4/5) | Haploid | |
|  | CF1379 blastospore 26 (44/57, 10 sites, site 7 4/5) |  | |
|  | CF1379 blastospore 27 (44/57, 10 sites, site 7 4/5) |  | |
|  | CF1379 blastospore 28 (44/57, 10 sites, site 8 5/5) | Haploid | |
|  | CF1379 blastospore 29 (44/57, 10 sites, site 8 5/5) | Haploid | |
|  | CF1379 blastospore 30 (44/57, 10 sites, site 8 5/5) |  | |
|  | CF1379 blastospore 31 (44/57, 10 sites, site 8 5/5) |  | |
|  | CF1379 blastospore 32 (44/57, 10 sites, site 8 5/5) |  | |
|  | CF1379 blastospore 33 (44/57, 10 sites, site 9 6/6) | Haploid | |
|  | CF1379 blastospore 34 (44/57, 10 sites, site 9 6/6) | Haploid | |
|  | CF1379 blastospore 35 (44/57, 10 sites, site 9 6/6) |  | |
|  | CF1379 blastospore 36 (44/57, 10 sites, site 9 6/6) |  | |
|  | CF1379 blastospore 37 (44/57, 10 sites, site 9 6/6) |  | |
|  | CF1379 blastospore 38 (44/57, 10 sites, site 9 6/6) |  | |
|  | CF1379 blastospore 39 (44/57, 10 sites, site 10 6/6) | Diploid | |
|  | CF1379 blastospore 40 (44/57, 10 sites, site 10 6/6) | Diploid | |
|  | CF1379 blastospore 41 (44/57, 10 sites, site 10 6/6) |  | |
|  | CF1379 blastospore 42 (44/57, 10 sites, site 10 6/6) |  | |
|  | CF1379 blastospore 43 (44/57, 10 sites, site 10 6/6) |  | |
|  | CF1379 blastospore 44 (44/57, 10 sites, site 10 6/6) |  | |
| *cks2*∆-2 | CF1387 blastospore 1 (40/53, 10 sites, site 1 4/4) | Haploid | |
|  | CF1387 blastospore 2 (40/53, 10 sites, site 1 4/4) | Haploid | |
|  | CF1387 blastospore 3 (40/53, 10 sites, site 1 4/4) |  | |
|  | CF1387 blastospore 4 (40/53, 10 sites, site 1 4/4) |  | |
|  | CF1387 blastospore 5 (40/53, 10 sites, site 3 3/4) | Haploid | |
|  | CF1387 blastospore 6 (40/53, 10 sites, site 3 3/4) | Haploid | |
|  | CF1387 blastospore 7 (40/53, 10 sites, site 3 3/4) |  | |
|  | CF1387 blastospore 8 (40/53, 10 sites, site 4 5/5) | Diploid | |
|  | CF1387 blastospore 9 (40/53, 10 sites, site 4 5/5) | Diploid | |
|  | CF1387 blastospore 10 (40/53, 10 sites, site 4 5/5) |  | |
|  | CF1387 blastospore 11 (40/53, 10 sites, site 4 5/5) |  | |
|  | CF1387 blastospore 12 (40/53, 10 sites, site 4 5/5) |  | |
|  | CF1387 blastospore 13 (40/53, 10 sites, site 5 5/6) | Haploid | |
|  | CF1387 blastospore 14 (40/53, 10 sites, site 5 5/6) | Haploid | |
|  | CF1387 blastospore 15 (40/53, 10 sites, site 5 5/6) |  | |
|  | CF1387 blastospore 16 (40/53, 10 sites, site 5 5/6) |  | |
|  | CF1387 blastospore 17 (40/53, 10 sites, site 5 5/6) |  | |
|  | CF1387 blastospore 18 (40/53, 10 sites, site 6 5/5) | Haploid | |
|  | CF1387 blastospore 19 (40/53, 10 sites, site 6 5/5) | Haploid | |
|  | CF1387 blastospore 20 (40/53, 10 sites, site 6 5/5) |  | |
|  | CF1387 blastospore 21 (40/53, 10 sites, site 6 5/5) |  | |
|  | CF1387 blastospore 22 (40/53, 10 sites, site 6 5/5) |  | |
|  | CF1387 blastospore 23 (40/53, 10 sites, site 7 5/5) | Diploid | |
|  | CF1387 blastospore 24 (40/53, 10 sites, site 7 5/5) | Diploid | |
|  | CF1387 blastospore 25 (40/53, 10 sites, site 7 5/5) |  | |
|  | CF1387 blastospore 26 (40/53, 10 sites, site 7 5/5) |  | |
|  | CF1387 blastospore 27 (40/53, 10 sites, site 7 5/5) |  | |
|  | CF1387 blastospore 28 (40/53, 10 sites, site 8 2/4) | Diploid | |
|  | CF1387 blastospore 29 (40/53, 10 sites, site 8 2/4) | Diploid | |
|  | CF1387 blastospore 30 (40/53, 10 sites, site 9 5/5) | Diploid | |
|  | CF1387 blastospore 31 (40/53, 10 sites, site 9 5/5) | Diploid | |
|  | CF1387 blastospore 32 (40/53, 10 sites, site 9 5/5) |  | |
|  | CF1387 blastospore 33 (40/53, 10 sites, site 9 5/5) |  | |
|  | CF1387 blastospore 34 (40/53, 10 sites, site 9 5/5) |  | |
|  | CF1387 blastospore 35 (40/53, 10 sites, site 10 6/8) | Diploid | |
|  | CF1387 blastospore 36 (40/53, 10 sites, site 10 6/8) | Diploid | |
|  | CF1387 blastospore 37 (40/53, 10 sites, site 10 6/8) |  | |
|  | CF1387 blastospore 38 (40/53, 10 sites, site 10 6/8) |  | |
|  | CF1387 blastospore 39 (40/53, 10 sites, site 10 6/8) |  | |
|  | CF1387 blastospore 40 (40/53, 10 sites, site 10 6/8) |  | |
| *P_GAL7_-CLB3* | CF1715 blastospore 1 (3/40, 15 sites, site1 2/3) germinated on YPG | Haploid | |
|  | CF1715 blastospore 2 (3/40, 15 sites, site1 2/3) germinated on YPG | Haploid | |
|  | CF1715 blastospore 3 (3/40, 15 sites, site10 1/4) germinated on YPG | Haploid | |
| **Plasmid** | **Genotype** | **Sources** | |
| pAG32 | *HYG AMP* | [4] | |
| pAI3 | *NAT AMP* | [5] | |
| pJAF1 | *NEO AMP* | [6] | |
| pJAF15 | *HYG AMP* | [6] | |
| pXL1 | *P_GPD1_* *NEO* *KAN AMP* | [7] | |
| pXL1-Cas9-HygB | *CAS9-HYGB* | [8] | |
| pYF515 | *sgRNA* scaffold | [9] | |
| pSDMA25 | *SH-NAT AMP* (*C. neoformans*) | [10] | |
| pCF3 | *SH-NEO AMP* (*C. deneoforman*) | This study | |
| pNURAT | *NURAT* in pAI3 | This study | |
| pCF7 | pNURAT in pCF3 | This study | |
| pSH5 | *P_GAL7_-CLB3-NEO KAN AMP* | This study | |

* Blastospore information (survival of all blastospores dissected for the given strain, total number of sites, the site that the blastospore is derived from, survival of blastospores dissected from this site).

** For each budding site, no more than two blastospores were chosen for FACS determination of ploidy.

1.	Lin, X., Hull, C.M., and Heitman, J. (2005). Sexual reproduction between partners of the same mating type in *Cryptococcus neoformans*. Nature *434*, 1017-1021.

2.	Zhai, B., Zhu, P., Foyle, D., Upadhyay, S., Idnurm, A., and Lin, X. (2013). Congenic strains of the filamentous form of *Cryptococcus neoformans* for studies of fungal morphogenesis and virulence. Infect. Immun. *81*, 2626-2637.

3.	Ni, M., Feretzaki, M., Li, W., Floyd-Averette, A., Mieczkowski, P., Dietrich, F.S., and Heitman, J. (2013). Unisexual and heterosexual meiotic reproduction generate aneuploidy and phenotypic diversity *de novo* in the yeast *Cryptococcus neoformans*. PLoS Biol. *11*, e1001653.

10.	Arras, S.D., Chitty, J.L., Blake, K.L., Schulz, B.L., and Fraser, J.A. (2015). A genomic safe haven for mutant complementation in *Cryptococcus neoformans*. PLoS ONE *10*, e0122916.
