## Supplementary material for "Dynamic genome plasticity during unisexual reproduction in the human fungal pathogen *Cryptococcus deneoformans*": Table S6.docx

**Table S6. Primers used in this study.**

| **Primer Name** | **Sequence (5' to 3')** | **Description** |
| --- | --- | --- |
| **Real-time PCR primers** | | |
| JOHE44120 | GTCTCCACTGATTTCATTGGCTCTAC | *GPD1* RT-F |
| JOHE44121 | GTAACCATACTCATTGTCATACCAGCTG | *GPD1* RT-R |
| JOHE42837 | AGCCCACACTCTCCTACTG | *KAR5* RT-F |
| JOHE42838 | TGTTCTTCGCTGCCACAT | *KAR5* RT-R |
| JOHE44867 | CAGCGTCTCCTGTCTCCTCT | CNA01720 RT-F |
| JOHE44868 | CCAGATACCCGCACATTTCT | CNA01720 RT-R |
| JOHE44869 | ACTCGGAAGGCCGTAACACT | CNA04030 RT-F |
| JOHE44870 | TGGAGGAGCAAACCTTGAAC | CNA04030 RT-R |
| JOHE44871 | GACTGCTGCTGAGGCAAATAC | CNA04230 RT-F |
| JOHE44872 | TCCCTTACTGCTCGCCTTCT | CNA04230 RT-R |
| JOHE44873 | CTACGATGAAGACGGCAAAC | CNA04250 RT-F |
| JOHE44874 | TAGCAGGTGCGAGAACAGAA | CNA04250 RT-R |
| JOHE44879 | GTTGAGCAAGGACGACAACA | CNC01750 RT-F |
| JOHE44880 | TCCAACGGACCACATATCAA | CNC01750 RT-R |
| JOHE44883 | ATGCATCCCTCCTGTCTCTC | CNE00250 RT-F |
| JOHE44884 | CCGTCAGAAGTATCGGGTGT | CNE00250 RT-R |
| JOHE44885 | GGCAGCAGTAGCGCATTAG | CNE02070 RT-F |
| JOHE44886 | CTTCCATGTCTTCCGTCAAG | CNE02070 RT-R |
| JOHE44887 | GGGTTCCACGACGAGATATT | CNE04100 RT-F |
| JOHE44888 | CTTCAACCCACCGTCTTACC | CNE04100 RT-R |
| JOHE44889 | TACGAACCAAGCCCTCAATC | CNE04400 RT-F |
| JOHE44890 | CCTCTTCCTCCATGTCATCCT | CNE04400 RT-R |
| JOHE44891 | CTTACACCGCCGAAGAGAAA | CNF01190 RT-F |
| JOHE44892 | CGACGGAAGAGGAGGACAT | CNF01190 RT-R |
| JOHE44893 | CCTGGAAAGCAACCAGCTAC | CNF02160 RT-F |
| JOHE44894 | CTATTGCCTCTTGGCGTTGG | CNF02160 RT-R |
| JOHE44895 | CAATCCAATGTGCAGGTGTC | CNG01990 RT-F |
| JOHE44896 | ATCCAAGGGAGGAGCAATGT | CNG01990 RT-R |
| JOHE44897 | GCTTTACGCACCAGCTTCTT | CNH00290 RT-F |
| JOHE44898 | GGCTGTTTGGCATGTTTAGC | CNH00290 RT-R |
| JOHE44899 | GTAACGAAGGCCGTTGAGAA | CNH01970 RT-F |
| JOHE44900 | AGACCAAAGTCGGCAATCA | CNH01970 RT-R |
| JOHE44901 | CGAAGTGGCTAGGAGGAACA | CNI01430 RT-F |
| JOHE44902 | TCGAACGACAGGCTTTGAG | CNI01430 RT-R |
| JOHE44903 | ATGGGTGTCAAGGATGGAGA | CNJ00160 RT-F |
| JOHE44904 | GAGTCGGAATTTGGAGTGGA | CNJ00160 RT-R |
| JOHE44905 | CACTGCCCTTCCCATCTTT | CNK01480 RT-F |
| JOHE44906 | CGCTGTGTGAGACCATAGTGA | CNK01480 RT-R |
| JOHE44909 | TACCATCGCCTAGCTTCACC | CNM00930 RT-F |
| JOHE44910 | TTTAACGACGCCTTGACTGG | CNM00930 RT-R |
| JOHE44911 | GAAAGGTCGAGGAGGTGATGT | CNM00990 RT-F |
| JOHE44912 | CGCAACCTTCTGATCCTCAT | CNM00990 RT-R |
| JOHE44913 | CAACTTATTGATCGCCGCTAA | CNN01700 RT-F |
| JOHE44914 | GGGCTGGATCAAATCTCAAA | CNN01700 RT-R |
| **Gene deletion primers** | | |
| M13F | GTAAAACGACGGCCAGT | *NAT*, *NEO*, and *HYG* cassettes |
| M13R | CAGGAAACAGCTATGAC | *NAT*, *NEO*, and *HYG* cassettes |
| **Deletion of *PCL2*** | | |
| JOHE45103 | GTCATAGCTGTTTCCTGgcatagaggaggatctcatagtg | CNA01720 3F |
| JOHE45104 | gcccgcaaacatctagtta | CNA01720 3R |
| JOHE45105 | tgcgcttgtcgtgaagttag | CNA01720 5F |
| JOHE45106 | ACTGGCCGTCGTTTTACcggagggatagcgagatgtat | CNA01720 5R |
| JOHE45109 | atttcacacgcagtcatcagg | CNA01720 Junc3R |
| JOHE45110 | acaaaccacgccacctatt | CNA01720 Junc5F |
| **Deletion of *PCL9*** | | |
| JOHE45111 | GTCATAGCTGTTTCCTGggtatttgggaactctgatgg | CNA04250 3F |
| JOHE45112 | attcttgggccttctacctct | CNA04250 3R |
| JOHE45113 | gaccgaagaccgaaacctaaa | CNA04250 5F |
| JOHE45114 | ACTGGCCGTCGTTTTACcagagaatgcaggcgaaagt | CNA04250 5R |
| JOHE45117 | tagatgaccacgaagatgacg | CNA04250 Junc3R |
| JOHE45118 | ccaacagcagcaataacagc | CNA04250 Junc5F |
| **Deletion of *PCL6*** | | |
| JOHE45143 | GTCATAGCTGTTTCCTGgtttgttctcaatcctcgca | CNK01480 3F |
| JOHE45144 | aagaccagagggcatattg | CNK01480 3R |
| JOHE45145 | ctcaagtaacaggttgcgttc | CNK01480 5F |
| JOHE45146 | ACTGGCCGTCGTTTTACttctcaaggatgtcagggatg | CNK01480 5R |
| JOHE45149 | aagggatgtatcaatgtggc | CNK01480 Junc3R |
| JOHE45150 | gcgcacgcttacgagattatt | CNK01480 Junc5F |
| **Deletion of *CKS1*** | | |
| JOHE45135 | GTCATAGCTGTTTCCTGgggtttcatttgttgagggtag | CNF02160 3F |
| JOHE45136 | aaagatggactggaatgtgc | CNF02160 3R |
| JOHE45137 | ttgagggaagtagtagtggatg | CNF02160 5F |
| JOHE45138 | ACTGGCCGTCGTTTTACcagtgaagttgagagaaagagg | CNF02160 5R |
| JOHE45141 | gacaagcattcgttgcagat | CNF02160 Junc3R |
| JOHE45142 | cattgtgtggagaatgatgg | CNF02160 Junc5F |
| **Deletion of *CKS2*** | | |
| JOHE45127 | GTCATAGCTGTTTCCTGccctctctcgcctttgtattg | CNF01190 3F |
| JOHE45128 | attacctcctcaccctcttgc | CNF01190 3R |
| JOHE45129 | gccgaccgatgcttaaata | CNF01190 5F |
| JOHE45130 | ACTGGCCGTCGTTTTACttgaagtggaggtctctgtca | CNF01190 5R |
| JOHE45133 | acaagttcggctctgatggt | CNF01190 Junc3R |
| JOHE45134 | gacgaacgagcgagatgatt | CNF01190 Junc5F |
| ***GAL7-CLB3* strain construction** | | |
| JOHE45941 | ggtgacgctgtgagagtgg | *CAS9*-F |
| JOHE45942 | gggcccctcttcacgtgg | *CAS9*-R |
| JOHE45943 | tttgcattagaactaaaaacaaagca | U6 F |
| JOHE45944 | TAAAACAAAAAAgcaccgactcggtgcc | gRNA R |
| JOHE46352 | gacttgaaattgcgaggttggttttagagctagaaatagcaag | CNE04400-sgRNA-F |
| JOHE46353 | caacctcgcaatttcaagtccaacagtataccctgccggtg | CNE04400-sgRNA-R |
| JOHE46478 | gcaccttcgttatcacatgcAACGTCGTGACTGGGAAAAC | pSH5-Plasmid backbone-F |
| JOHE46479 | catagccccaatcttcgTGTGAAATTGTTATCCGCTC | pSH5-Plasmid backbone-R |
| JOHE45726 | cgaagattggggctatg | pSH5-CNE04400 5UTR-F |
| JOHE46348 | ACTGGCCGTCGTTTTACagggtagtggcgagtgagg | pSH5-CNE04400 5UTR-NEO-R |
| JOHE46337 | GTCATAGCTGTTTCCTGgggcagtacaggctaagcgt | pSH5-NEO-PGAL7-F |
| JOHE46338 | ggtaactcgagtttgttcaaaaa | pSH5-PGAL7-R |
| JOHE46351 | tttttgaacaaactcgagttaccatggcttcccgagtaagtatcc | pSH5-PGAL7-CNE04400-F |
| JOHE46350 | gcatgtgataacgaaggtgc | pSH5-CNE04400-R |
| JOHE45301 | ggacgtttaggtcacgttagaa | NEO-GAL7-CNE04400-F |
| JOHE46452 | tccatgtagcgagggttagg | NEO-GAL7-CNE04400-R |
| **NURAT strain construction** | | |
| JOHE42262 | GACTCGAAGTCTTTGGGCAGATCTGATGAGGTGGCACT | pCF3-Pasmid backbone-F |
| JOHE42267 | CCAGCTCACATCCTCGCAGCCATGGACTCAACCCTATCTC | pCF3-Pasmid backbone-R |
| JOHE42268 | GAGATAGGGTTGAGTCCATGGCTGCGAGGATGTGAGCTGG | pCF3-NEO-F |
| JOHE42265 | TTGGACGTGCTTTATTGGCCGGTTTATCTGTATTAACACGGAAG | pCF3-NEO-R |
| JOHE42263 | TTAATTAACACAAGTATCGTGGCGCGCCCACAAGATGTTTTACATCAGTA | pCF3-SH-5' homology-F |
| JOHE42261 | AGTGCCACCTCATCAGATCTGCCCAAAGACTTCGAGTC | pCF3-SH-5' homology-R |
| JOHE42264 | GGCGCGCCACGATACTTGTGTTAATTAAACCAAACTAATCTGTGTCAAAA | pCF3-SH-3' homology-F |
| JOHE42266 | CTTCCGTGTTAATACAGATAAACCGGCCAATAAAGCACGTCCAA | pCF3-SH-3' homology-R |
| JOHE40981 | CAGCTGAAGCTTCGTACGC | JM41 HYG-F (pAG32) |
| JOHE40982 | GCATAGGCCACTAGTGGATCTG | JM42 HYG-R (pAG32) |
| JOHE41858 | CAGATCCACTAGTGGCCTATGCGGACACAAGATGATGTCGAAG | URA5 3F |
| JOHE41546 | TAAGTCATTCCATCCCCTCG | URA5 3R |
| JOHE41543 | CCAGGACGCATATTGCTTCC | URA5 5F |
| JOHE41857 | GCGTACGAAGCTTCAGCTGGTCTTGCTTCAGAGACAGTG | URA5 5R |
| JOHE42010 | CAGTAGTACAGCCATCAGGC | URA5 Junc3R |
| JOHE42009 | CCTGGTCGCTCAAACTATCG | URA5 Junc5F |
| JOHE41353 | CATGGTCATAGCTGTTTCCTG | pNURAT Plasmid backbone-F (pAI3) |
| JOHE41352 | ATTCACTGGCCGTCGTTTTAC | pNURAT Plasmid backbone-R (pAI3) |
| M13F | GTAAAACGACGGCCAGT | pNURAT-M13F for NAT-5' |
| JOHE41548 | GAGCTTGCTCTCCGTCAGATG | pNURAT-NATCDS+580R |
| JOHE41549 | CATCTGACGGAGAGCAAGCTCGCAAAGAGCGAAGTTGCTCG | pNURAT-URA5-F |
| JOHE41550 | CTGTAACGGTAAGCCGTATCGGTTCAGAGTTTGATTGGACGA | pNURAT-URA5-R |
| JOHE41547 | CGATACGGCTTACCGTTACAG | pNURAT-NATCDS+21F |
| M13R | CAGGAAACAGCTATGAC | pNURAT-M13R for NAT-3' |
| M13F | GTAAAACGACGGCCAGT | pCF7 NURAT-F (pNURAT) |
| M13R | CAGGAAACAGCTATGAC | pCF7 NURAT-R (pNURAT) |
| JOHE41353 | CATGGTCATAGCTGTTTCCTG | pCF7 Plasmid backbone-F (pCF3) |
| JOHE41352 | ATTCACTGGCCGTCGTTTTAC | pCF7 Plasmid backbone-R (pCF3) |
| JOHE42274 | CAGATGCCCTGGATGATGTC | SH Junc5F |
| JOHE42278 | CTTAGTTCGGACGACCTCCAG | SH Junc3R |
| JOHE41444 | CGGACGAGCTCTCAAATTGG | Sxi2_Da_Forward |
| JOHE41445 | TTTGCTCGCTCTCCTTCCAC | Sxi2_Da_Reverse |
| JOHE41446 | GCCGTGCAAGGGTGTAGG | Sxi1_Dalpha_Foward |
| JOHE41447 | GGGCCATTGGAGGAAGCTG | Sxi1_Dalpha_Reverse |
| **Chromoblot probes** | | |
| JOHE41547 | CGATACGGCTTACCGTTACAG | NAT Probe-F |
| JOHE41548 | GAGCTTGCTCTCCGTCAGATG | NAT Probe-R |
| JOHE41861 | CGCTTCACAGTCCAATCGAA | URA5 Probe-F |
| JOHE41767 | TGCCAGAGGTAAGAACATCG | URA5 Probe-R |
| JOHE50076 | GCCACCTTGGTCACAGATAG | Chr2 segmental aneuploidy probe-F |
| JOHE50077 | GGCCCTTCCAgtcagtaca | Chr2 segmental aneuploidy probe-R |
| JOHE50084 | ACCATGTCCTCGCTCTCACTA | Chr6 segmental aneuploidy probe-F |
| JOHE50085 | CTCTTCCCTCCTCTCCTTGATAC | Chr6 segmental aneuploidy probe-R |
| JOHE50090 | tctgttgccgctgctttac | Chr13 segmental aneuploidy probe-F |
| JOHE50091 | TCAGGAGTTGCCGGTCATA | Chr13 segmental aneuploidy probe-R |
