## Supplementary figures and images for "Dynamic genome plasticity during unisexual reproduction in the human fungal pathogen *Cryptococcus deneoformans*"

### Figure S1.tif

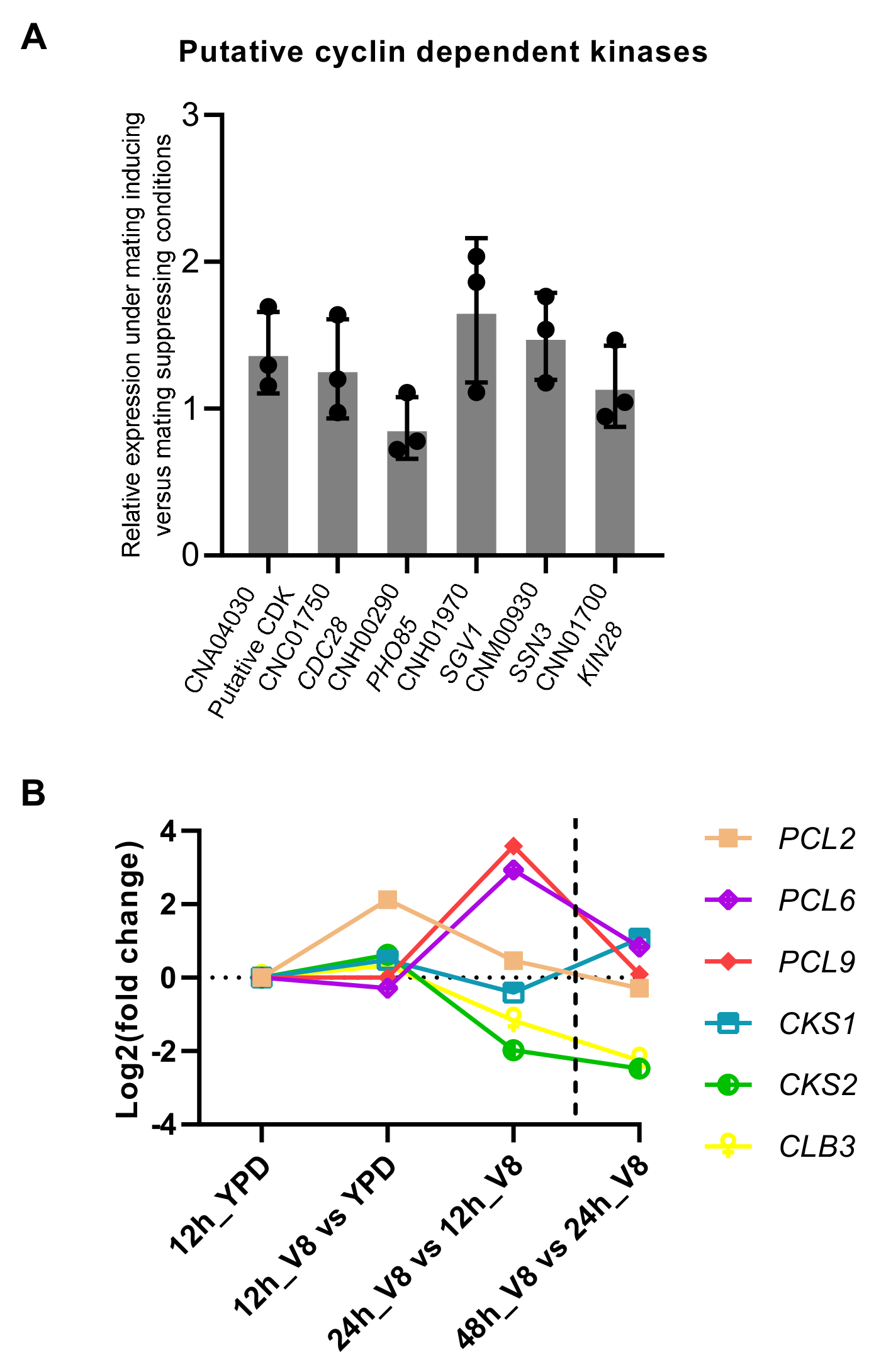

### Figure S2.tif

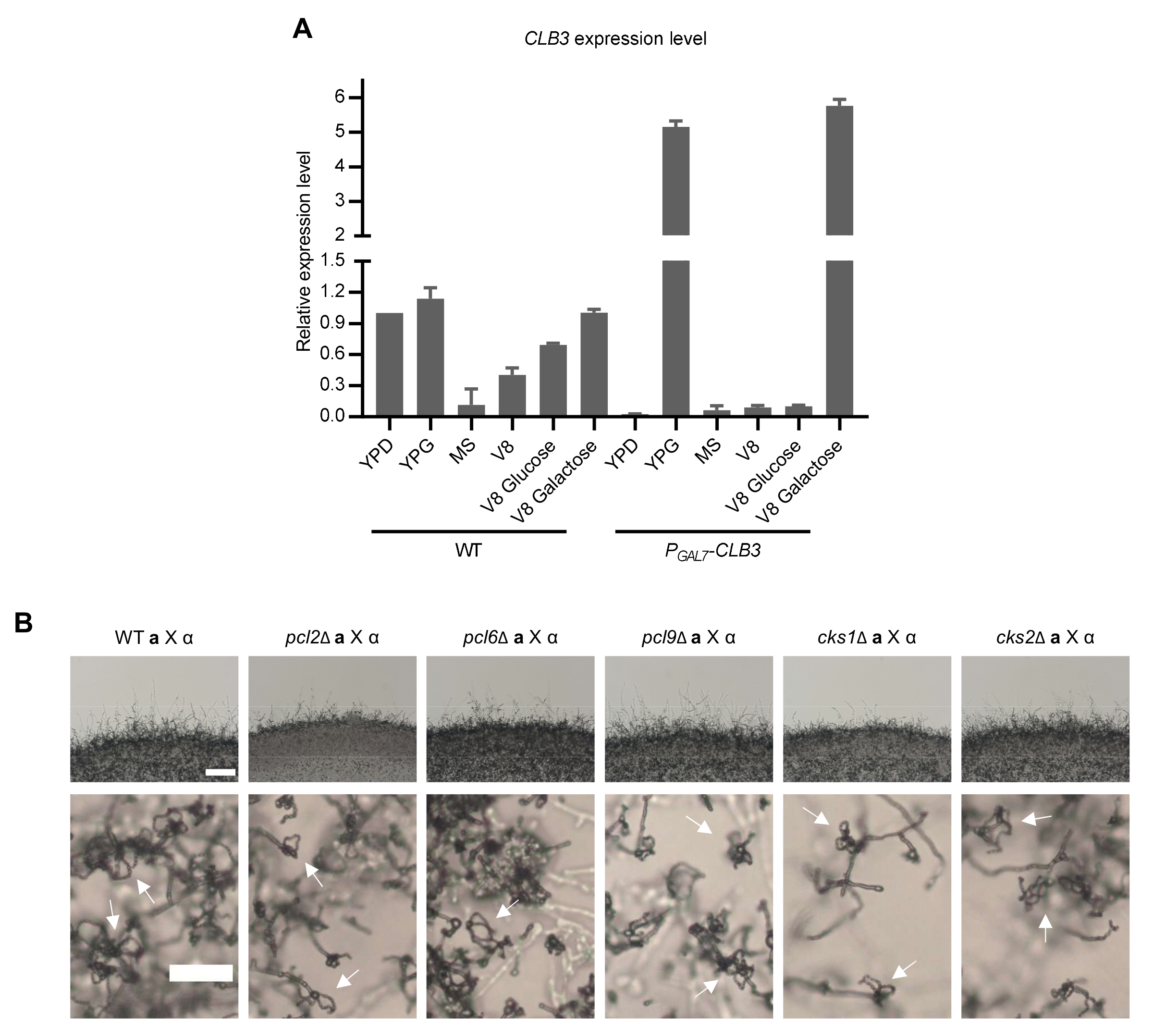

### Figure S3.tif

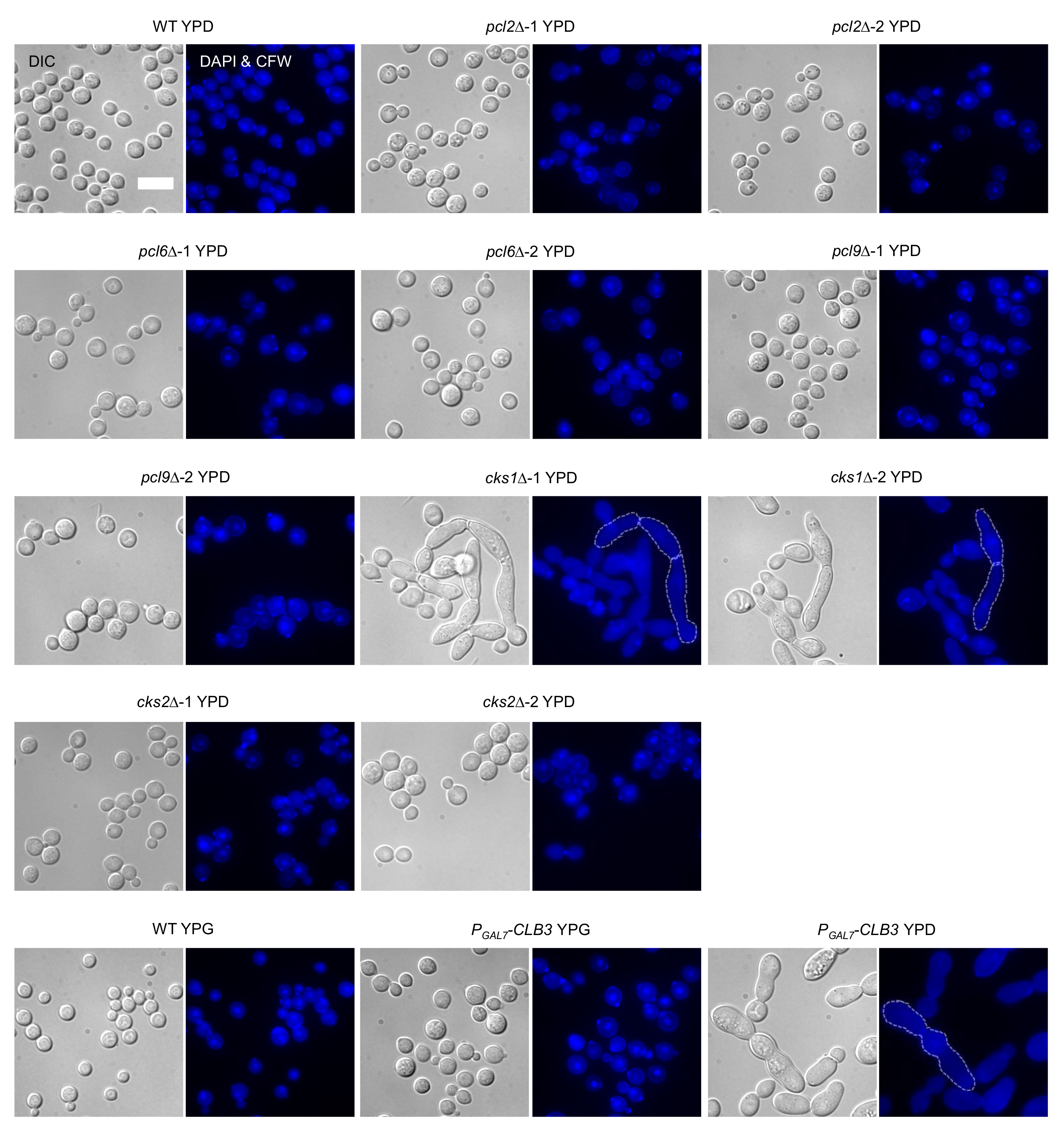

### Figure S4.tif

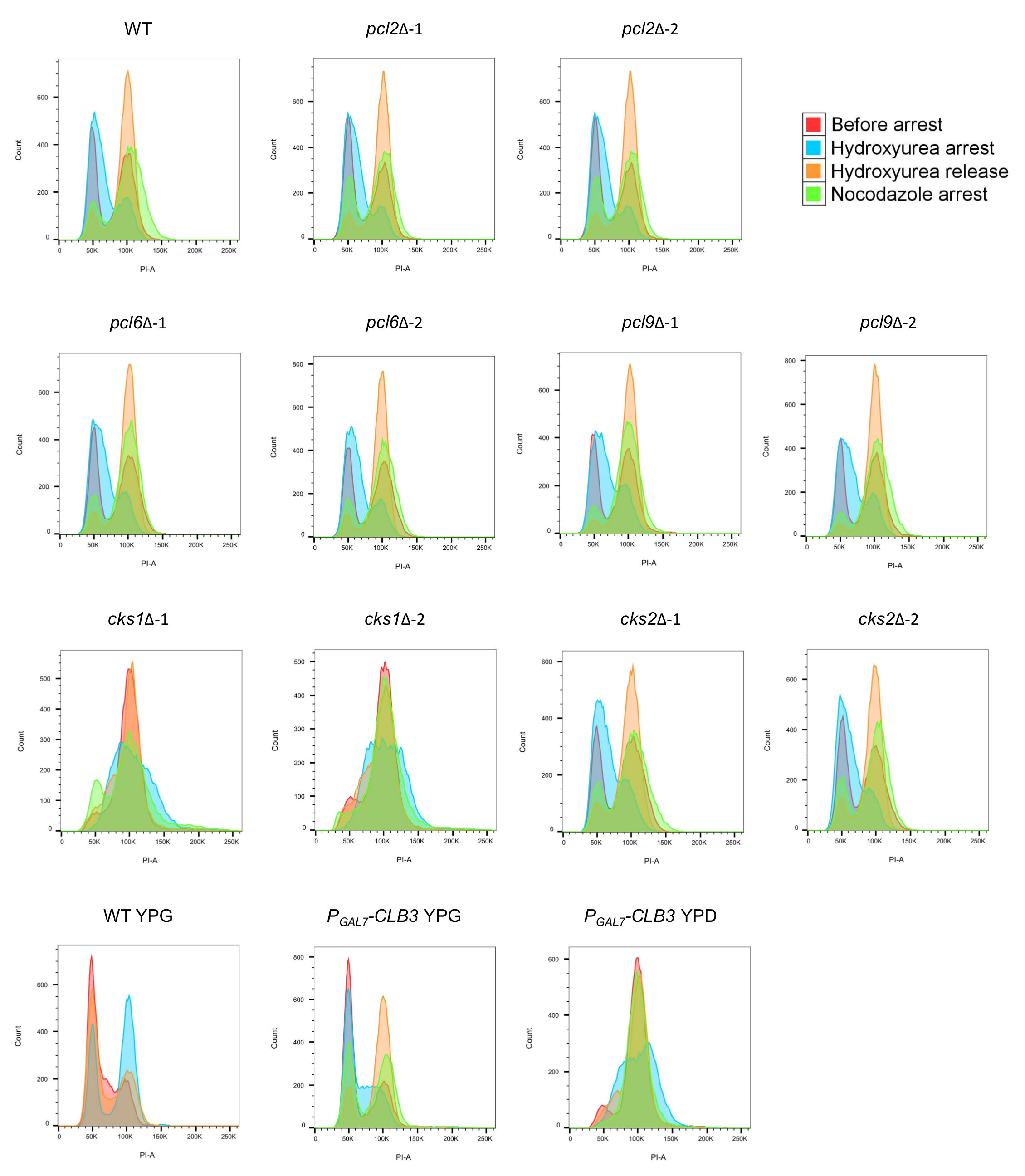

### Figure S5.tif

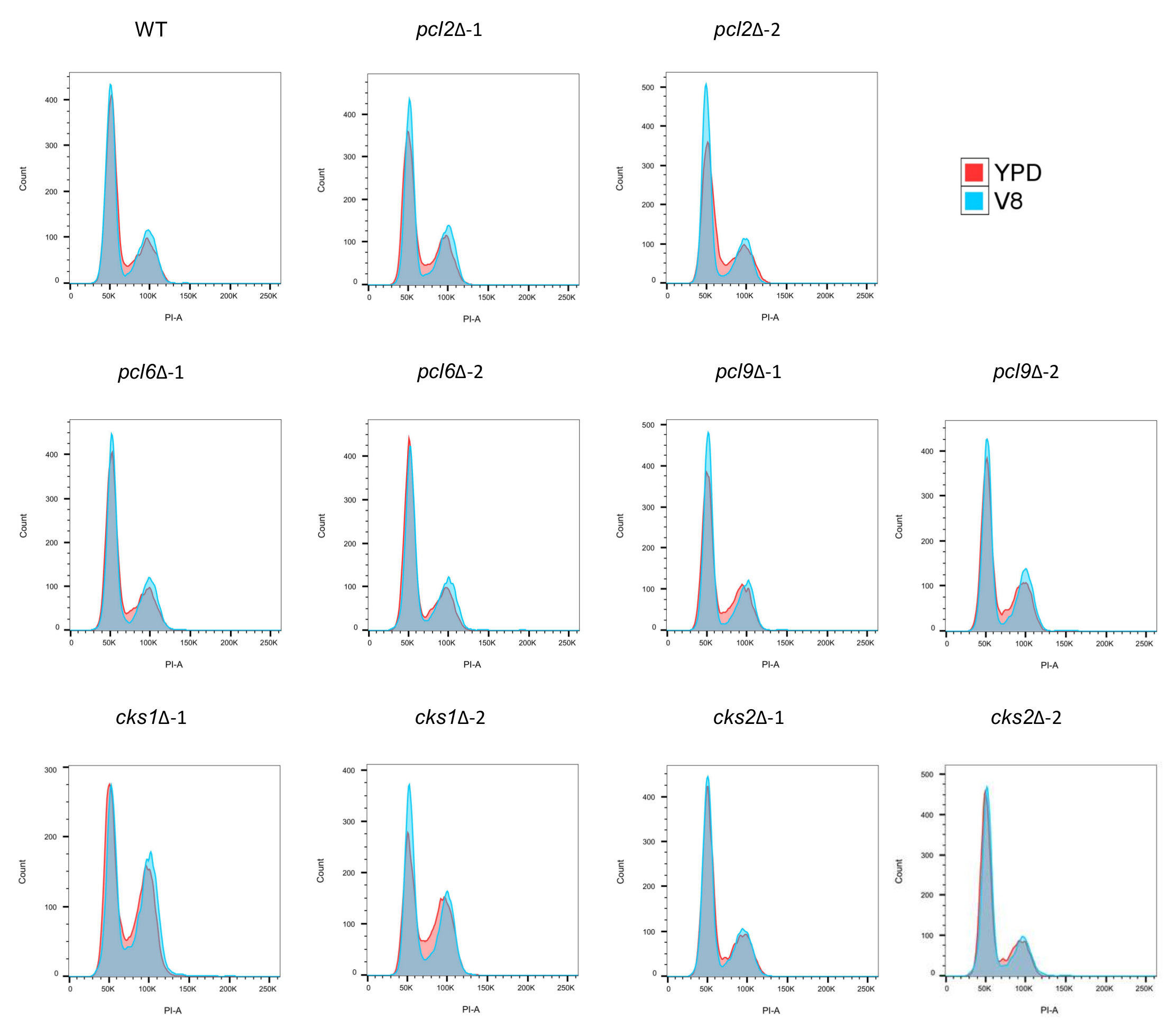

### Figure S6.tif

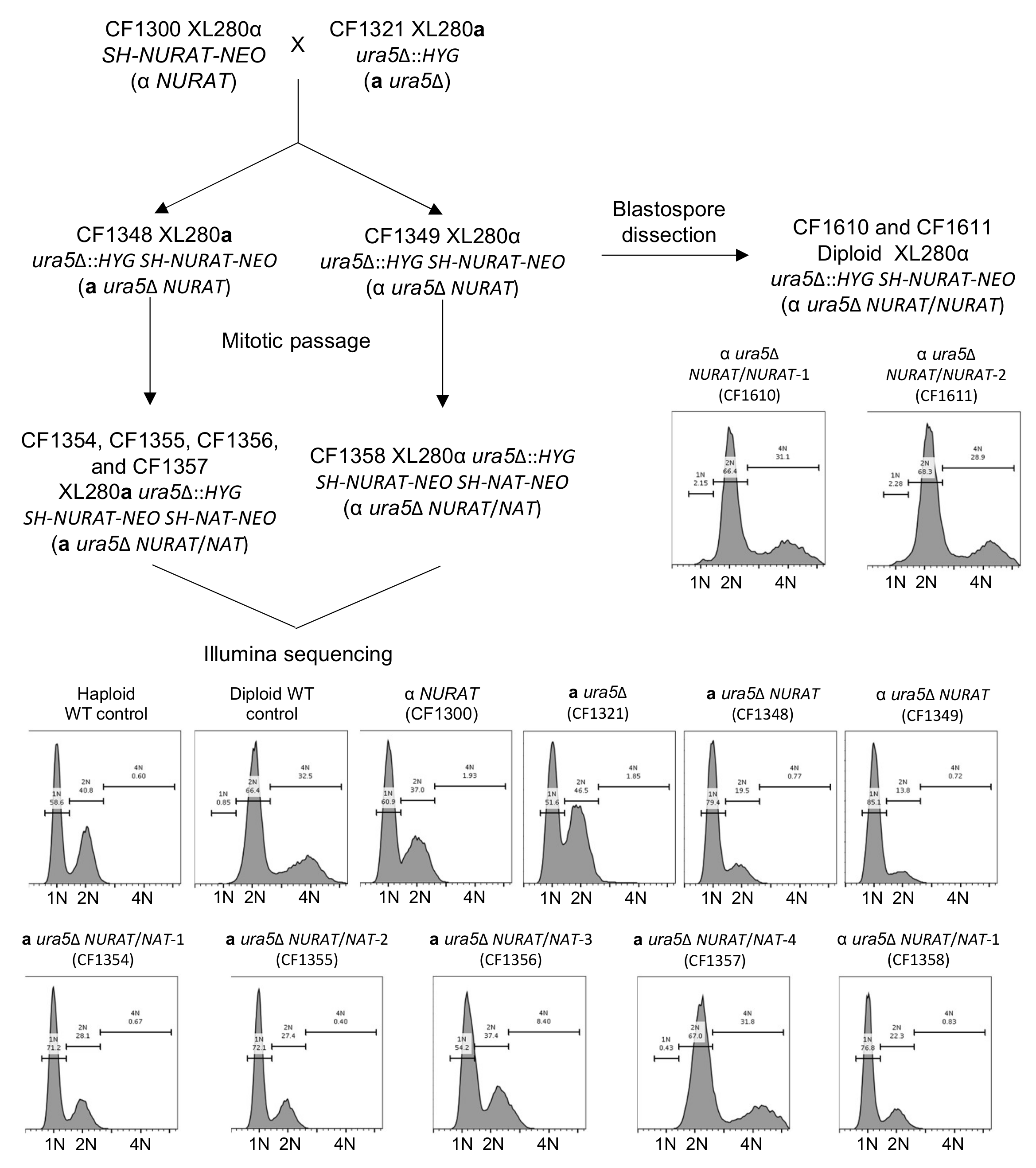

### Figure S7.tif

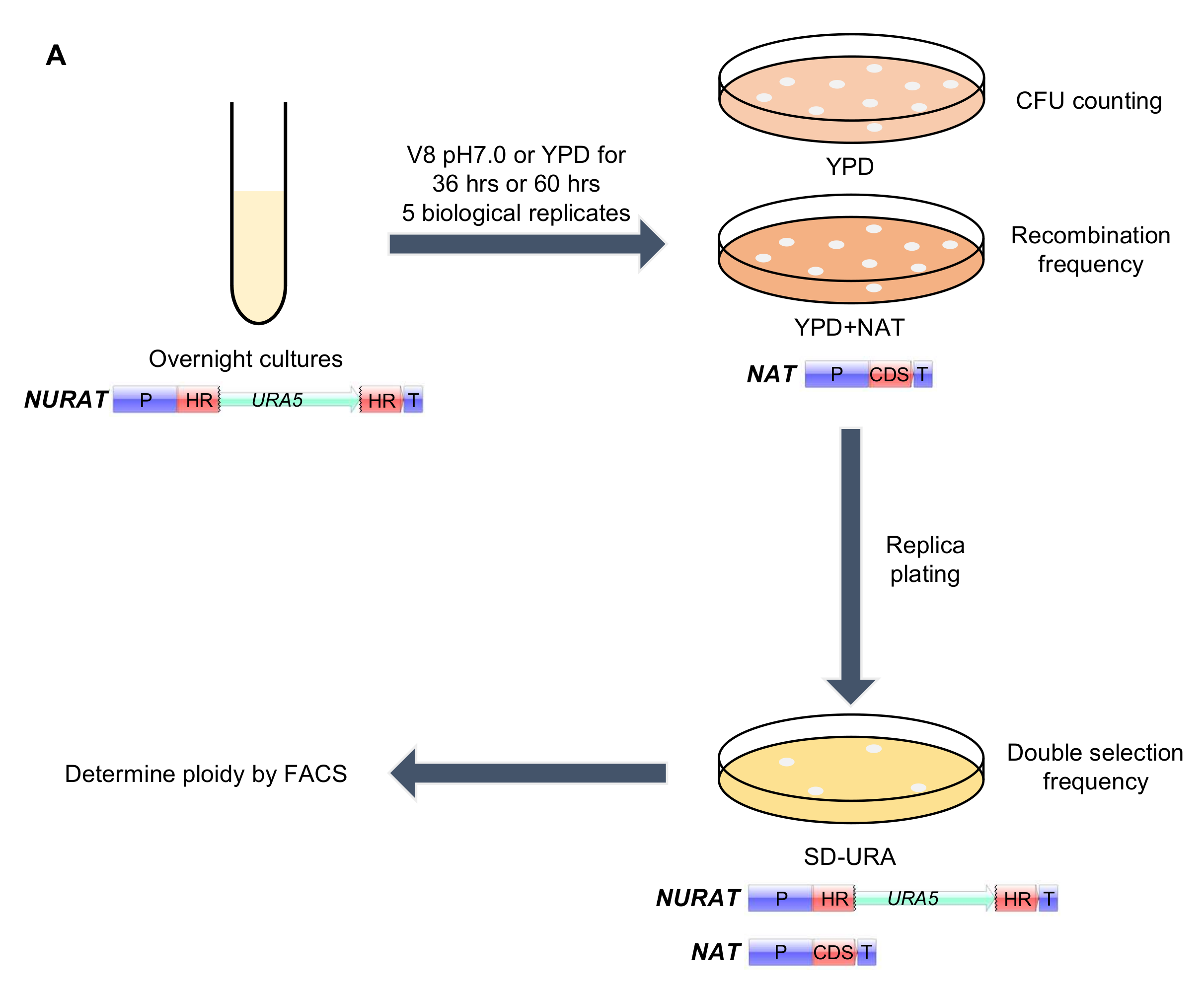

### Figure S8.tif

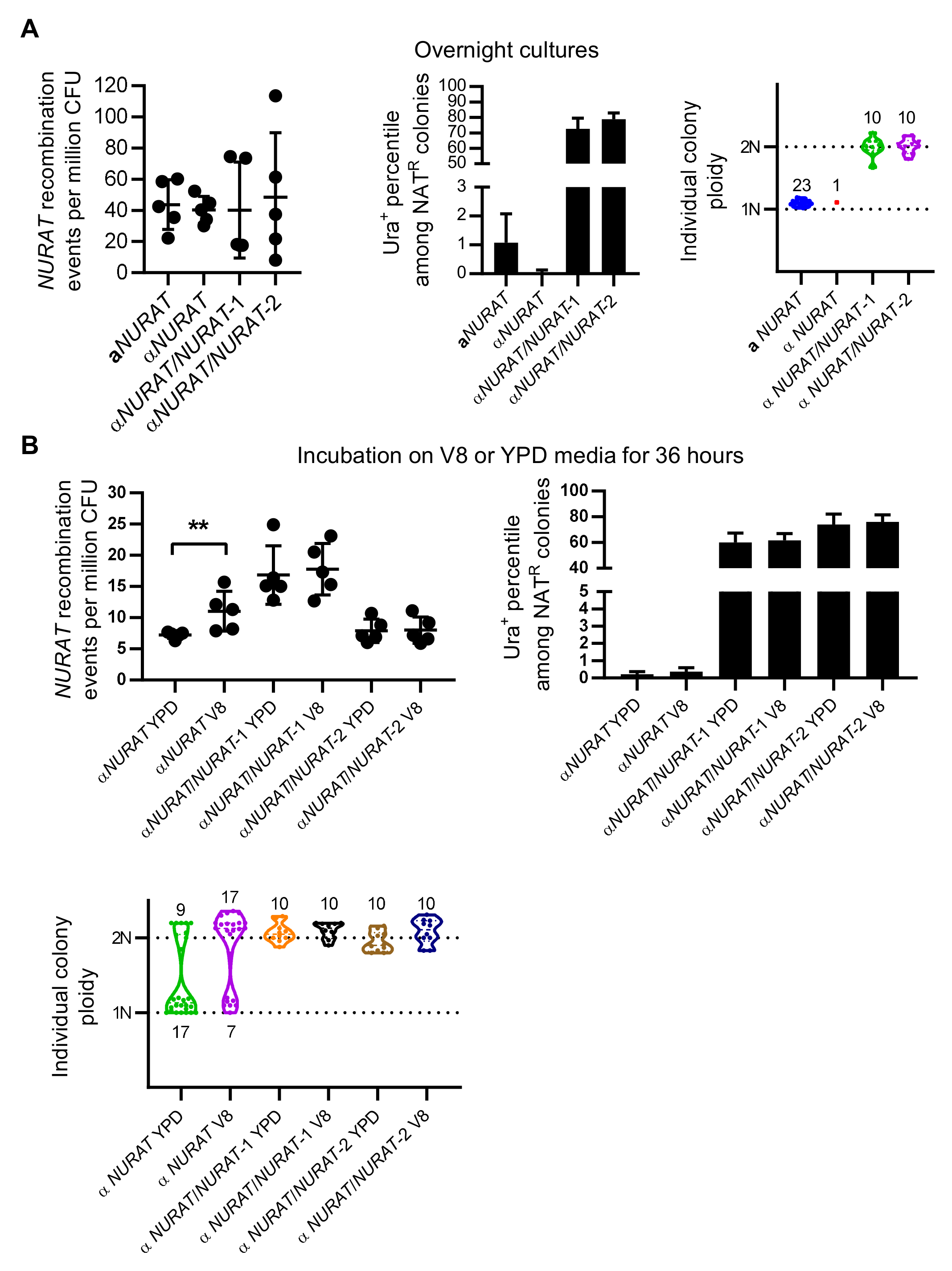

### Figure S9.tif

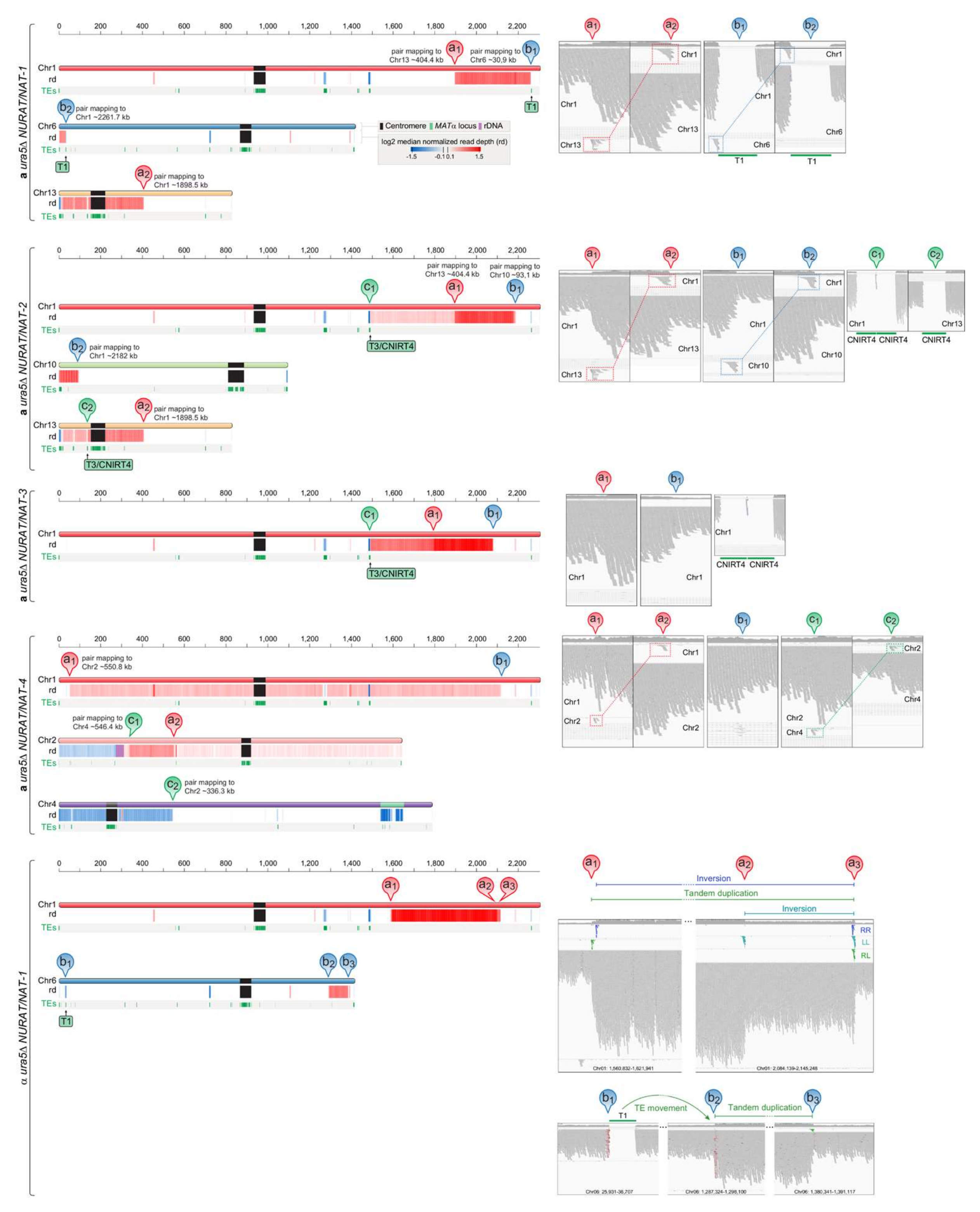

### Figure S10A-B.tif

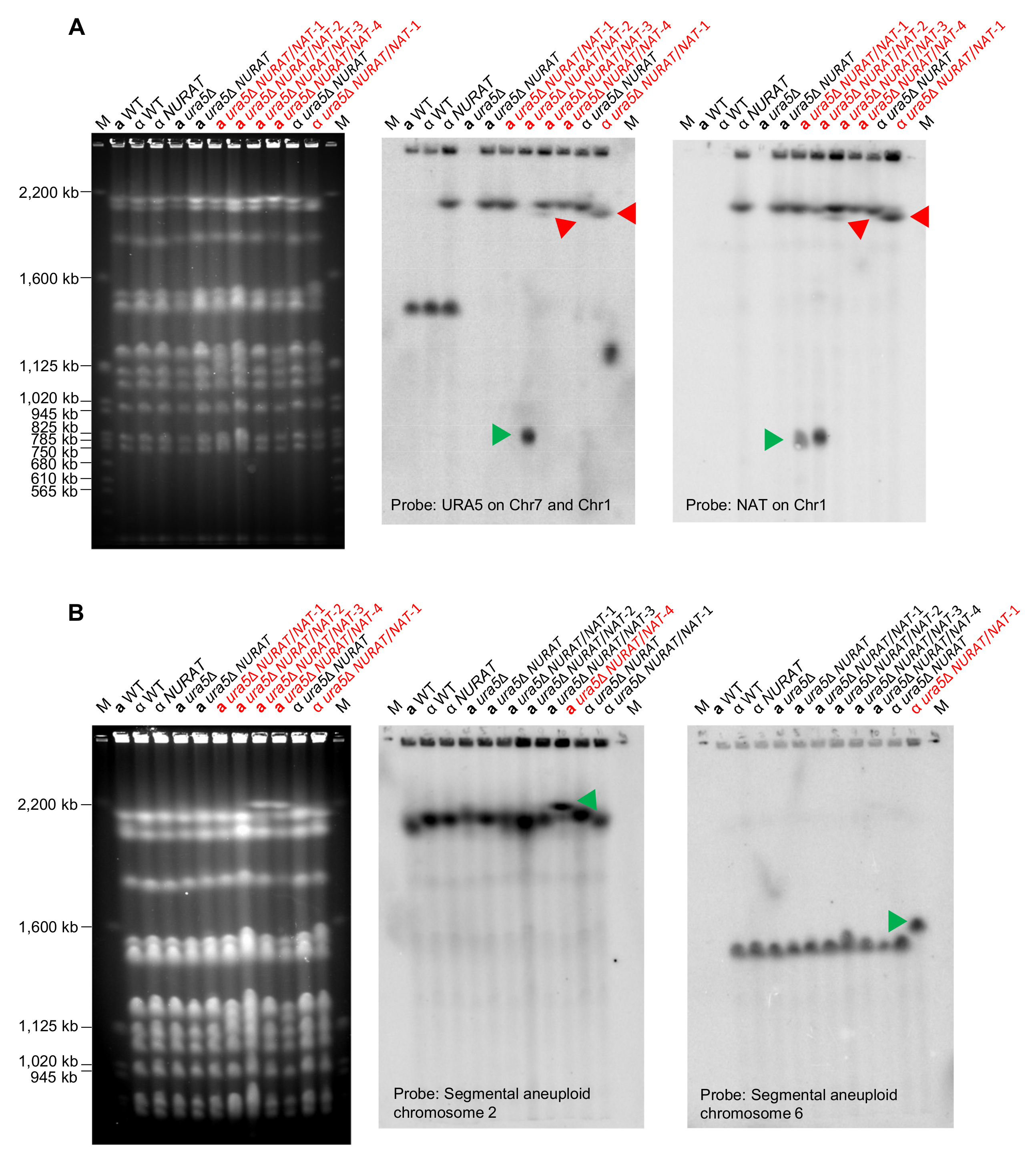

### Figure S10C.tif

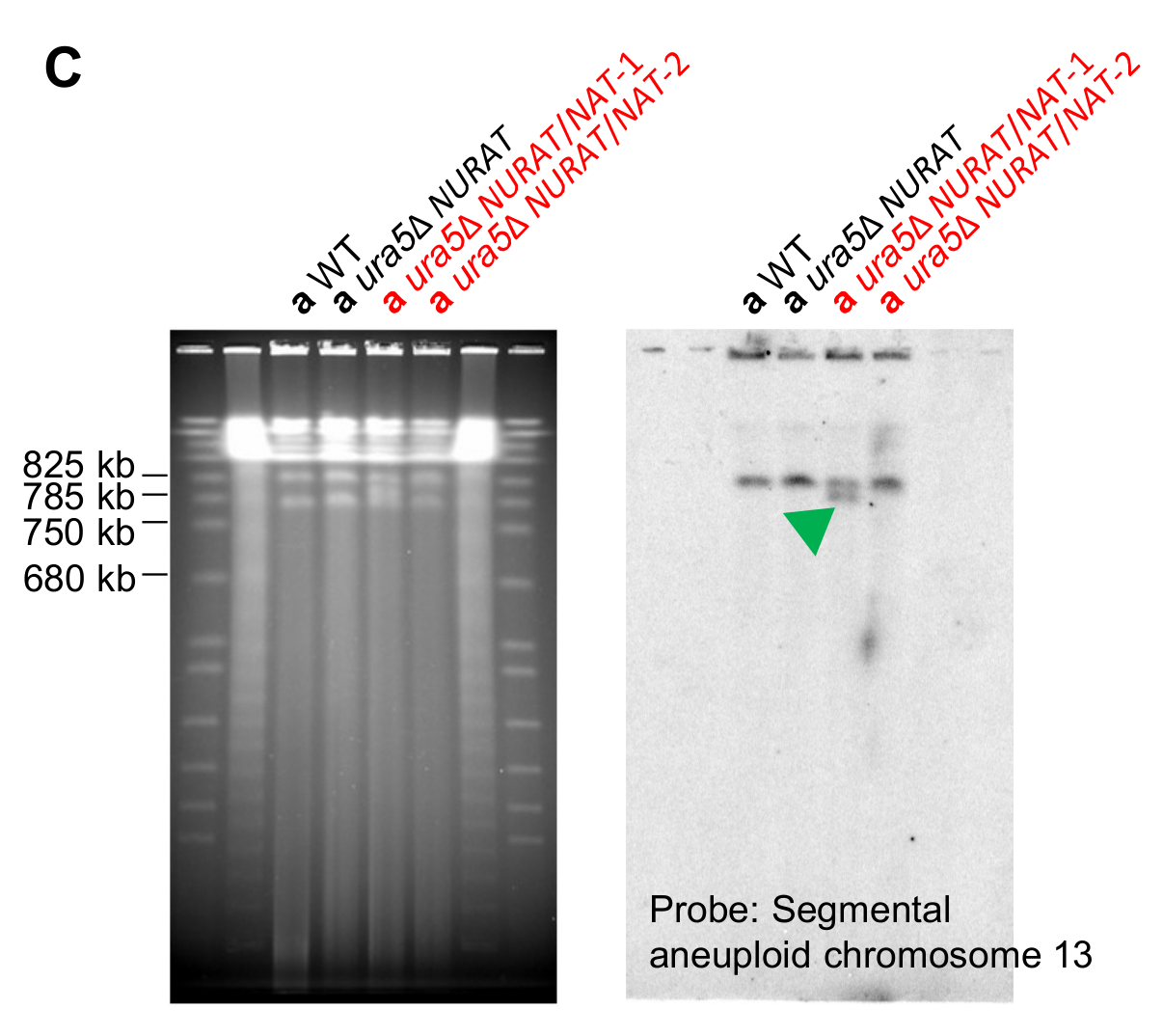

### Figure S11.tif

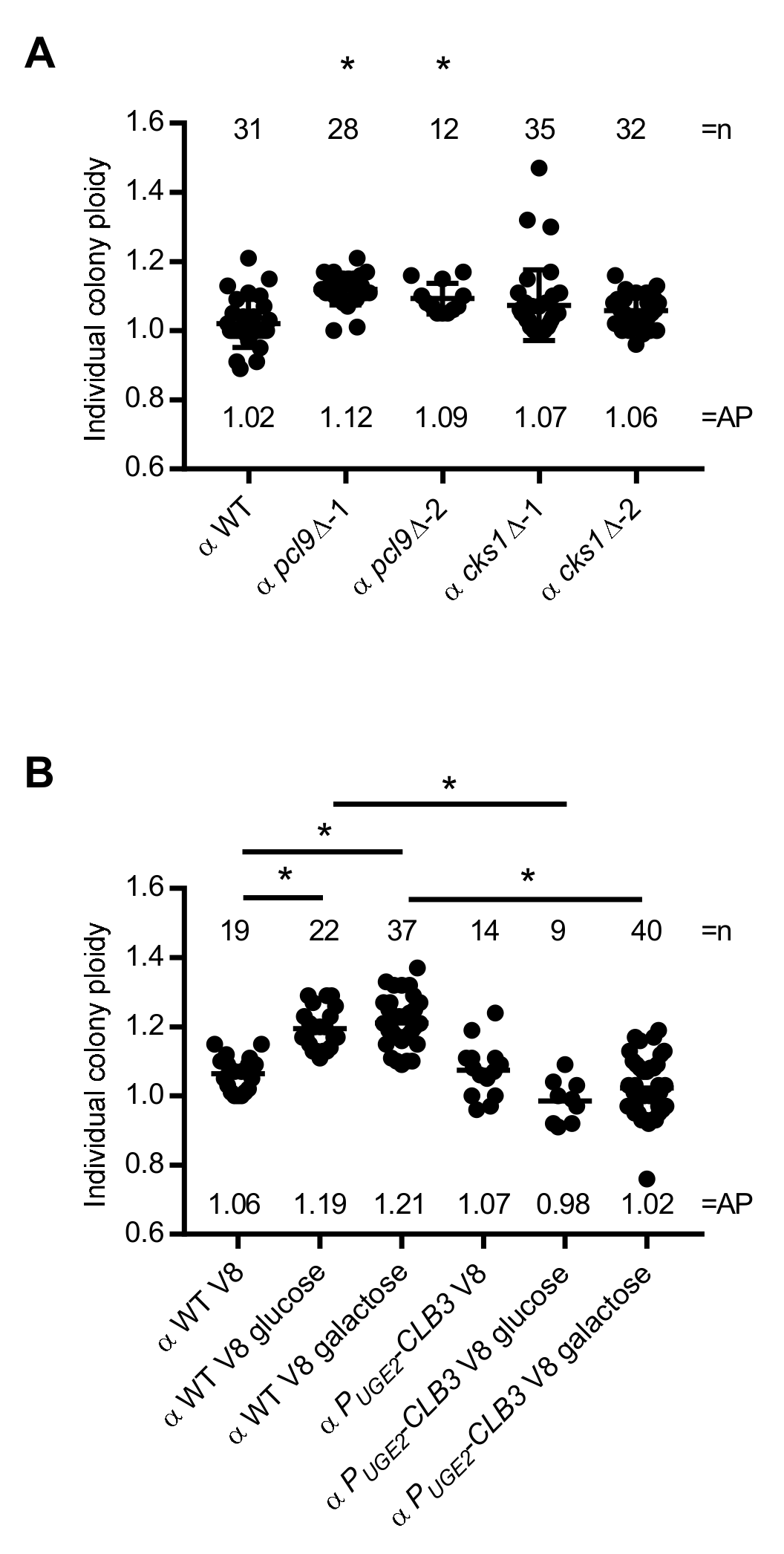
